# Visual Perception of 3D Space and Shape in Time - Part I: 2D Space Perception by 2D Linear Translation

**DOI:** 10.1101/2022.03.01.482161

**Authors:** Umaima Afifa, Javier Carmona, Amy Dinh, Diego Espino, Trevor McCarthy, Brian Ta, Patrick Wilson, Benjamin Asdell, Jinwoo Baik, Archana Biju, Sonia Chung, Christopher Dao, Mark Diamond, Saba Doust, Angela East, Diego Espino, Kailey Fleiszig-Evans, Adrian Franco, Anthony Garibay-Gutierrez, Aparajeeta Guha, Roshan Gunturu, Luke Handley, Christina Honore, Abinav Kannan, Jared Khoo, Mira Khosla, Chandan Kittur, Alexandra Kwon, Jessica Lee, Nicholas Lwe, Mylan Mayer, Elizabeth Mills, Delilah Pineda, Pasha Pourebrahim, Jacob Rajacich, Shan Rizvi, Liliana Rosales, Leonard Schummer, Conor Sefkow, Alexander Stangel, Cindy Ta, Ivy Ta, Natalie Tong, Kyle Tsujimoto, Alyssa Vu, Henry Wang, Amanda Yares, Natsuko Yamaguchi, Ki Woong Yoon, Shuyi Yu, Aaron P. Blaisdell, Katsushi Arisaka

## Abstract

Visual perception plays a critical role in navigating space and extracting useful semantic information crucial to survival. To identify distant landmarks, we constantly shift gaze vectors through saccades, while still maintaining the visual perception of stable allocentric space. How can we sustain stable allocentric space so effortlessly? To solve this question, we have developed a new concept of **NHT** (Neural Holography Tomography). This model states that retinotopy is invisible (not available to consciousness) and must be converted to a time code by traveling alpha brainwaves to perceive objects consciously. According to this framework, if identical alpha phases are continually assigned to a landmark, we perceive its exact and consistent allocentric location.

To test this hypothesis, we designed reaction time (RT) experiments to observe evidence of the predicted space-to-time conversion. Various visual stimuli were generated at a wide range of eccentricities either on a large TV (up to 40°) or by LED strips on a hemispherical dome (up to 60°). Participants were instructed to report the observed patterns promptly under either covert (no eye movement) or overt (with eye movement) conditions. As predicted, stimuli presented at the center of fixation always produced the fastest RTs. The additional RT delay was precisely proportional to the eccentricity of the peripheral stimulus presentation. Furthermore, both covert and overt attention protocols created the same RT delays, and trajectories of saccadic eye motions were in parallel to the overt RT vs. eccentricity. These findings strongly support our NHT model, in which the observed RT-eccentricity dependence is indicative of the spatiotemporal conversion required for maintaining a stable allocentric frame of reference. That is, we perceive space by time.

## 1 Introduction

Navigation of space is a complex task that we seem to be able to achieve effortlessly and, in most cases, unconsciously. To understand external space, we take a visual signal from the outer environment and convert it to a neural signal in the brain to reconstruct the external world, internally. While animals and humans use a variety of reference frames for spatial perception and memory (Trullier et al., 1997), the two main classes are: egocentric and allocentric. Using the egocentric strategy, animals use meaningful landmarks (local focus) and internally generated signals (e.g., vestibular, path integration, etc.) to orient in and navigate through 3D environments. An egocentric strategy allows the individual to learn the locations of specific targets and plan future movements through known space. In contrast, an allocentric strategy makes use of mental spatial maps where the subject orients itself according to distal landmarks (Jordan et al., 2004). Understanding absolute external space using an allocentric reference frame has important advantages as it is more flexible (O’Keefe & Nadel, 1978; but see Bouchekioua et al., 2021).

An allocentric strategy provides a critical advantage over an egocentric strategy in providing a means to anchor external objects to a spatial framework independent of the individual’s point of view. Anchoring objects in allocentric space not only allows for their use to guide navigation through space, but also to allow for semantic processes of the objects, such as their identity and category membership. Recognizing objects from various distances, orientations, and perspectives is important to be able to determine prey from predator,friend from foe, and to recognize the faces of specific individuals. The neural mechanisms by which objects are encoded in allocentric space and semantic information is bound to them are still unknown, despite significant efforts to understand both processes (Schneegans & Bays, 2019; Wang et al., 2021). A theory has been recently advanced that attempts to unite both processes through a single, coherent neural mechanism (Arisaka, 2022a, b, c). According to this theory, retinotopically encoded spatial information —which is necessarily egocentric—is scanned via alpha brain waves. The travelling brain wave, such as an alpha wave, takes a predictable amount of time to scan the retinotopic image from left to right and top to bottom, given the constant velocity of the brainwave. As a result, the time it takes the traveling wave to complete a scan from beginning to end of an object’s 2D retinotopic image in brain tissue allows for the temporal difference to be computed and stored. Thus, the compression of 2D spatial information into 1D temporal information using the principle of Neuro Holographic Tomography (NHT) allows the brain to convert spatial information into a time code. The compression of space into a hologram allows for semantic information, such as color, shape, and meaning, to be efficiently bound together. Arisaka (2022a, b) proposes that a Holographic Lattice Ring Attractor (HAL) provides the universal neural mechanism for storing semantic object information including its bound location in external allocentric space.Once an object and its properties are bound, they can be unpacked in consciousness for recognition and action.For example, by encoding the features of a face in a holographic representation, any face we observe can then be compared to our memory of faces to determine if the face we see belongs to someone we know or to a stranger.

This mechanism of holographic encoding is consistent with two processes of visual perception: unconscious and conscious. It has been well established, on the one hand, that unconscious processing of sudden visual onset happens rapidly (200 ms) and in parallel such that a sudden onset anywhere in the visual field can be detected equipotentially. Such rapid detection and reaction to sudden onset stimuli occurs prior to conscious processing. Conscious visual perception, on the other hand, typically takes about 400 ms to complete. More-over, semantic processing in conscious perception takes longer the further is the object from the center of vision (See Arisaka & Blaisdell, 2022 for review of unconscious and conscious visual perception).

We hypothesize that spatial information is encoded in the frequency time domain corresponding to shifts in the allocentric frame of reference. Thus, our theory uniquely predicts that reaction time (RT) to a visual onset will be independent of its eccentricity (opening angle from the center field of view) when no semantic information is necessary for a response, whereas RT will increase with eccentricity when responding is based on conscious processing of semantic information, such as in making a choice.

Previous experiments have tested the effect of eccentricity on various visual stimuli (Ando et al., 2016; Arkin & Yehuda, 1985; Bayle et al., 2011; Berlucchi et al., 1971; Marzi et al., 2006; Osaka, 1976; Rains, 1963; Schiefer et al., 2001) with all reporting higher reaction time for stimulus presented at the periphery. However,in these experiments the maximum eccentricity tested was 35 degrees (Berlucchi et al., 1971) under covert attention conditions only. Additionally, each experiment either tested simple reaction time (SRT)(Ando et al., 2016; Arkin & Yehuda, 1985; Berlucchi et al., 1971; Marzi et al., 2006; Osaka, 1976; Rains, 1963; Schiefer et al., 2001) or choice reaction time (CRT) (Bayle et al., 2011). There is a gap in experimental study comparing different attention conditions at larger peripheral locations testing semantic and non-semantic information processing.

The present study aims to fill this gap through a systematic study that tests visual stimulus response rate at various peripheral locations, up to 60 degrees along the right horizontal meridian (RHM) and the left horizontal meridian (LHM) and 40 degrees along the upper vertical meridian (UVM) and the lower vertical meridian (LVM). The behavioral tests were conducted under different attention conditions-without an overt fixation shift, that is a covert shift of attention and with a fixation-shift paradigm that is an overt shift in attention. Additionally, the overt attention condition tests oculomotor response through remote eye tracking. Subjects either overtly shifted their attention from a central location to a peripheral target by making an eye movement towards it, or they covertly shift their attention towards a peripheral target, while maintaining central fixation, in both cases making a manual response corresponding to the nature of the target. The study also tested how RT dependence on eccentricity compared in SRT and CRT and different level of complexity of semantic information.

## 2 Results

The objective of our study is to understand how spatial information is encoded in the brain where higher order processes are responsible for perception of visual stimulus. We postulate that semantic spatial information is converted to temporal information and developed simple reaction time and choice reaction time experiments to understand the internal mechanism. We test the visual stimuli processing along the horizontal meridian in Experiment 1(a, b, c), Experiment 2(a, c), and Experiment 3(a, b, c), and along the vertical meridian in Experiment 2b.

### 2.1 Reaction Time vs Eccentricity along Horizontal and Vertical axis

In the experiments, the participants performed reaction-based tasks where they were asked to respond as quickly as possible to target visual stimuli under different attention conditions, overt attention and covert attention (see additional details in Materials and Methods). The participants were instructed to press corresponding buttons to register the stimuli they observed. The stimuli were randomly flashed at different eccentricities along the horizontal or vertical meridian, depending on the protocol.

In Experiment 1, there were 6 protocols (SRT, 2-CRT and 3-CRT) for each stimulus set-gabor patterns (1a), English alphabetical letters (1b), and unfamiliar faces (1c). For SRT protocols, the participants were instructed to respond to a single stimulus that randomly appeared at different eccentricities along the horizontal meridian. For 2-CRT protocols, the subjects were randomly presented with one of two target stimuli at different eccentricities and had to select which of the two they observed. For 3-CRT protocols, the subjects were presented with one of three target stimuli. For the Gabor and Unfamiliar Face experiments, stimuli were presented at 0, 5, 10, 15, 20, 25, 30, 35, and 40 degrees away from the center; for the Familiar and Unfamiliar Character experiments, stimuli were presented at 0, 10, 20 30, and 40 degrees away from the center on both, left and right side.

In Experiment 2a, there were 4 protocols (SRT and 3-CRT) for stimulus set-BBB. The participants were instructed to press corresponding buttons to register the number of adjacent blue LEDs (1, 2 or 3) they observed. The LEDs were randomly flashed at different eccentricities along the horizontally aligned LED strip at 0-, 15-, 30-, 45-, or 60-degrees eccentricities from the center on both, left and right side. In experiment 2b, all four protocols discussed in Experiment 2a were repeated along the vertical meridian. All parameters of the experiment, except the eccentricity range, was same as the protocol along the horizontal axis. In experiment 2b, the LEDs were flashed along the vertically aligned LED strip at 0, 10, 20, 30 and 40 degrees along UVM and LVM. Experiment 2c was identical to experiment 1c.

In Experiment 3, there were 6 protocols (SRT, 2-CRT and 3-CRT) for each stimulus set-BBB (3a) and color (3b).For the BBB experiment, the experiment was identical to Experiment 2a. For the color experiment, the stimuli presented was either blue, red, or yellow LED. For the LR Experiment (3c), the subject was presented with a blue LED stimulus at the center and at the periphery with some time delay between the two and was instructed to respond whether they observed the stimuli at the center or the periphery first. The LEDs were randomly flashed at different eccentricities along the horizontally aligned LED strip at 0-, 15-, 30-, and 45-degrees eccentricities from the center on both, left and right side.

The average (mean) reaction time at each eccentricity is plotted along with the error bars (showing standard errors of the mean) for each data point. The average value excludes outliers as discussed in methods section. Figure 1 shows the eccentricity dependence of reaction time for all three experiments and the protocols, along the horizontal and vertical meridian aggregated over all participants. Example plots from experiment 2c is plotted in Figure 2 and 3 to show the mean RT vs eccentricity for each protocol for all subjects and the mean RT vs eccentricity and normalized mean RT vs eccentricity resepectively averaged over all subject (N=40). Key parameters, such as slope, intercept at 0 degree eccentricity, correlation coefficient of slopes and reduced *χ*^2^ values of the aggregated data is summarized in Table 1, 2 and 3. The slope and intercept data are also summarized in Figure 4.

**Figure (1).**
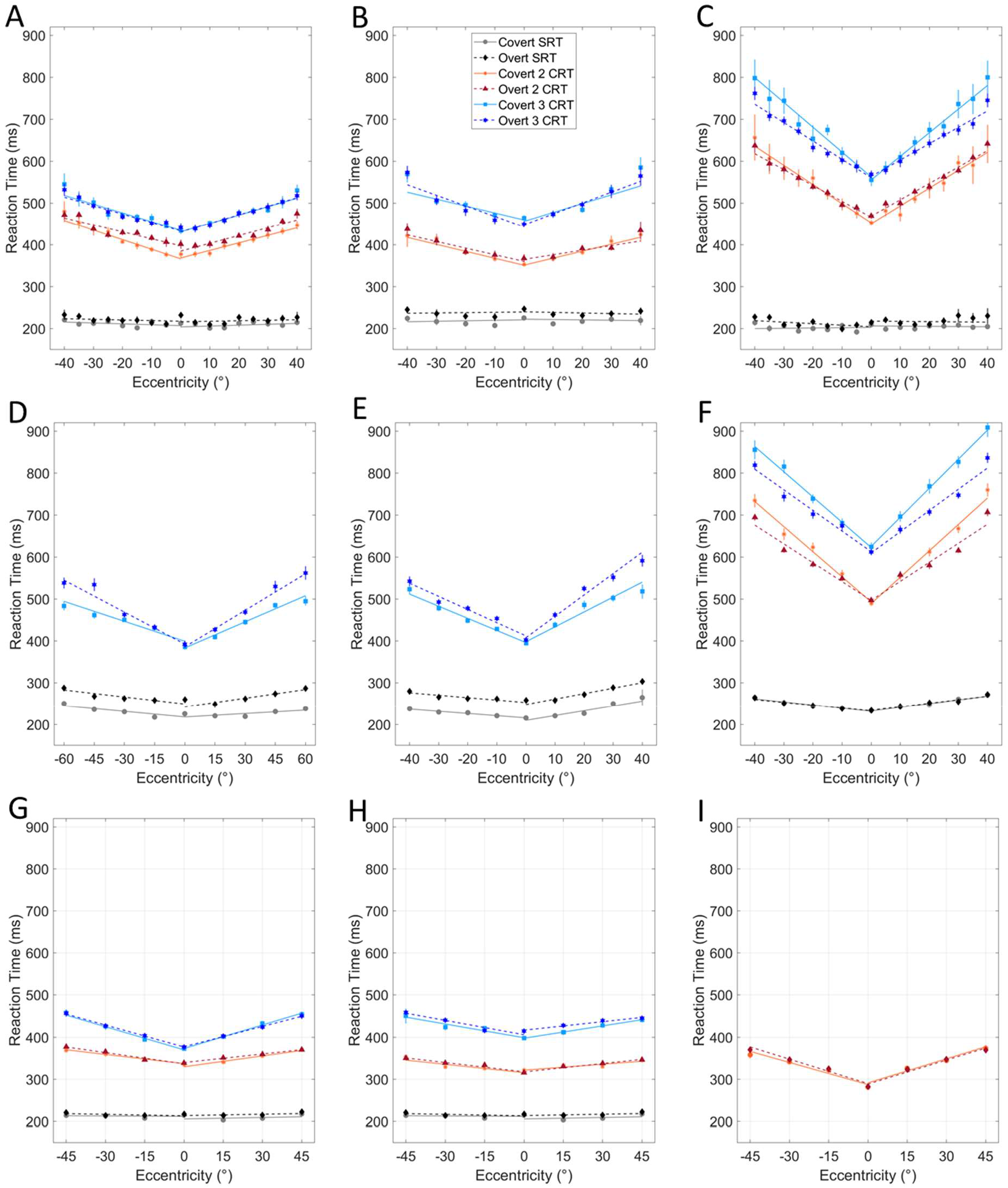
(A, B, C) Mean RT versus eccentricity of stimuli for Experiment 1a, 1b, 1c respectively. The protocols were plotted for aggregated data over all participants (N= 13). (D, E, F) Average RT versus eccentricity of stimuli for Experiment 2a, 2b, 2c. (D, E) The protocols were plotted for aggregated data over all participants (N= 13). ( F) The protocols were plotted for aggregated data over all participants (N= 40). (G, H, I) Mean reaction time versus eccentricity of stimuli for Experiment 3a, 3b, 3c. The protocols were plotted for aggregated data over all participants (N= 15). A best fit line using least-squares linear regression was plotted for both the positive eccentricities along the Horizontal Meridian (RHM) and Vertical Meridian (UVM) and negative eccentricities along the Horizontal Meridian(LHM) and Vertical Meridian (LVM) with error bars for each Mean RT.

**Figure (2).**
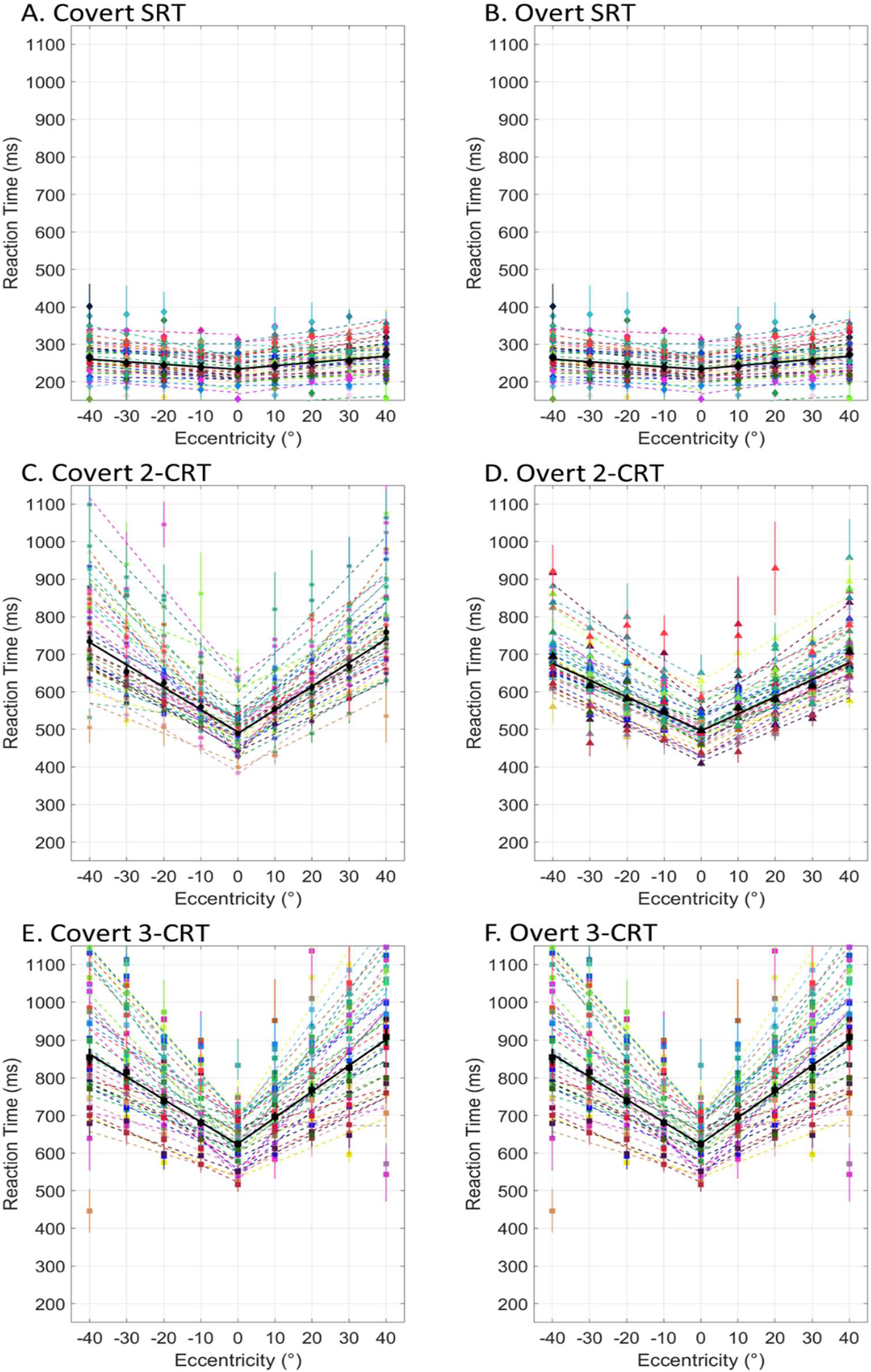
Mean reaction time versus distance of stimuli for all participants (N=40) for Experiment 2c. (A) Experiment 2c Covert SRT Protocol. (B) Experiment 2c Overt SRT Protocol. (C) Experiment 2c Covert 2-CRT Protocol. (D) Experiment 2c Overt 2-CRT Protocol. (E) Experiment 2c Covert 3-CRT Protocol. (F) Experiment 2c Overt 3-CRT Protocol.

**Figure (3).**
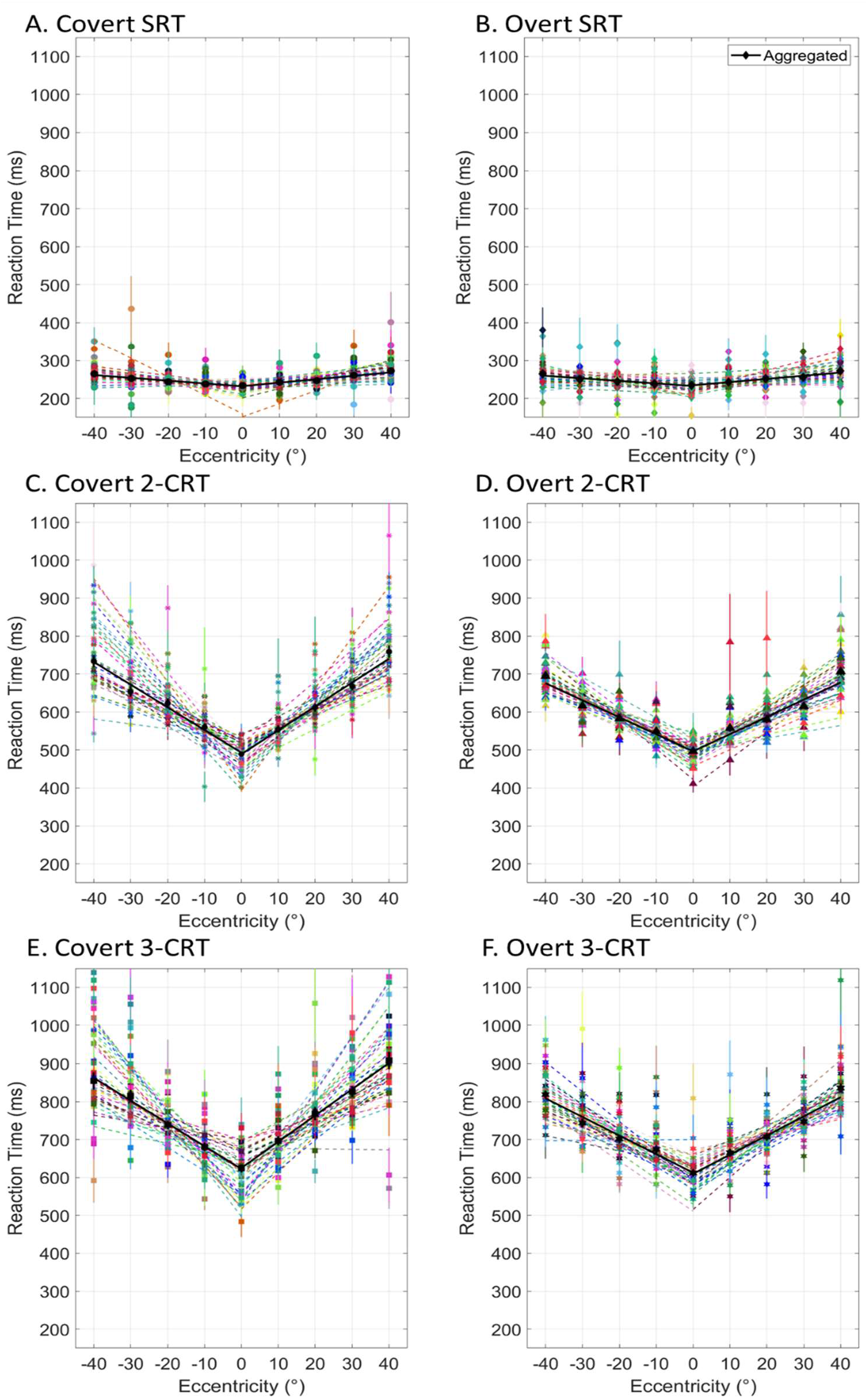
Mean normalized reaction time versus distance of stimuli for all participants (N=40) for Experiment 2c. (A) Experiment 2c Covert SRT Protocol. (B) Experiment 2c Overt SRT Protocol. (C) Experiment 2c Covert 2-CRT Protocol. (D) Experiment 2c Overt 2-CRT Protocol. (E) Experiment 2c Covert 3-CRT Protocol. (F) Experiment 2c Overt 3-CRT Protocol.

**Figure (4).**
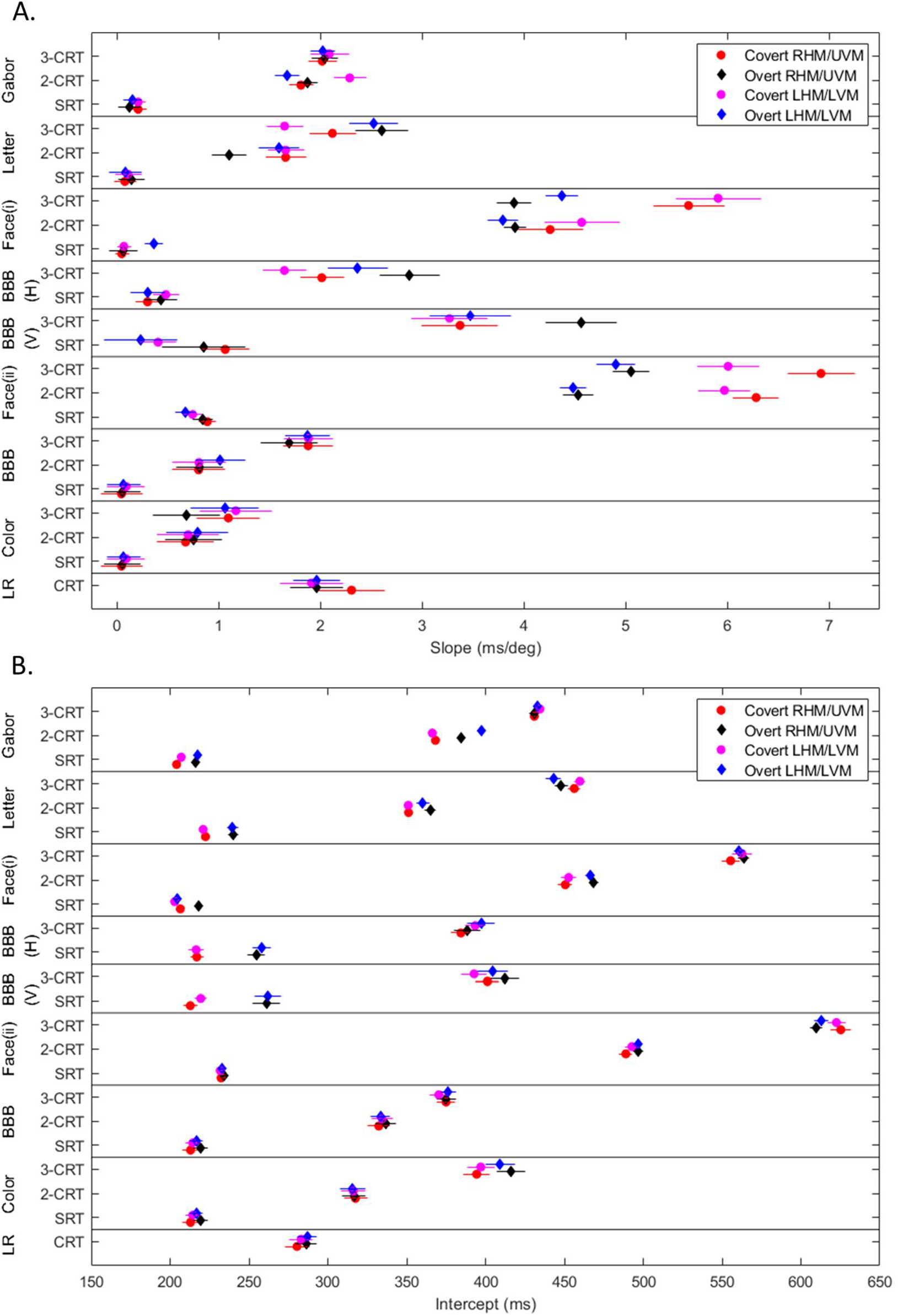
Slope and intercept data extracted through Chi Square minimization and the corresponding error bars are plot-ted for each protocol subdivided into the experiments. (A) Slope data for all protocols across all experiments conducted forRT to stimuli along Horizontal and Vertical Meridian. (B) Intercept data for all protocols across all experiments conducted for RT to stimuli along Horizontal and Vertical Meridian.

**Table (1).** Correlation coefficient *R*^2^, and reduced *χ*^2^ for the slope and intercept of regression lines in Figure 1(A, B, C). The data are aggregated over all participants (N= 13), categorized by protocol and direction of scanning.

|  | 0, -5, -10, -15, -20, -25, -30, -35, -40 (LHM) |  |  |  | 0, 5, 10, 15, 20, 25, 30, 35, 40 (RHM) |  |  |  |
| --- | --- | --- | --- | --- | --- | --- | --- | --- |
| Protocols | Slope (ms/°) | Intercept (ms) | $\chi^2_v$ | R | Slope (ms/°) | Intercept (ms) | $\chi^2_v$ | R |
| Covert SRT | 0.21 ± 0.07 | 207 ± 2 | 0.96 | 0.35 | 0.21 ± 0.08 | 204 ± 2 | 1.16 | 0.29 |
| Overt SRT | 0.15 ± 0.09 | 218 ± 2 | 1.26 | 0.40 | 0.12 ± 0.11 | 216 ± 2 | 1.74 | 0.21 |
| Covert 2-CRT | 2.29 ± 0.16 | 366 ± 3 | 0.57 | 0.98 | 1.81 ± 0.12 | 368 ± 3 | 0.33 | 0.98 |
| Overt 2-CRT | 1.67 ± 0.12 | 398 ± 3 | 1.66 | 0.93 | 1.87 ± 0.10 | 385 ± 2 | 1.12 | 0.94 |
| Covert 3-CRT | 2.09 ± 0.19 | 435 ± 3 | 0.53 | 0.96 | 2.02 ± 0.14 | 431 ± 3 | 0.54 | 0.98 |
| Overt 3-CRT | 2.02 ± 0.12 | 433 ± 3 | 1.07 | 0.96 | 2.04 ± 0.13 | 431 ± 3 | 0.30 | 0.99 |

Experiment 1a(Gabor) fit parameter data
|  | 0, -5, -10, -15, -20, -25, -30, -35, -40 (LHM) |  |  |  | 0, 5, 10, 15, 20, 25, 30, 35, 40 (RHM) |  |  |  |
| --- | --- | --- | --- | --- | --- | --- | --- | --- |
| Protocols | Slope (ms/°) | Intercept (ms) | $\chi^2_v$ | R | Slope (ms/°) | Intercept (ms) | $\chi^2_v$ | R |
| Covert SRT | 0.11 ± 0.13 | 221 ± 3 | 2.60 | 0.15 | -0.08 ± 0.11 | 223 ± 2 | 1.34 | -0.04 |
| Overt SRT | 0.08 ± 0.16 | 240 ± 4 | 1.59 | 0.07 | -0.14 ± 0.13 | 240 ± 3 | 1.07 | -0.20 |
| Covert 2-CRT | 1.66 ± 0.18 | 351 ± 3 | 0.73 | 0.98 | 1.66 ± 0.20 | 351 ± 3 | 0.29 | 0.99 |
| Overt 2-CRT | 1.59 ± 0.20 | 360 ± 4 | 1.02 | 0.96 | 1.10 ± 0.17 | 365 ± 4 | 1.01 | 0.92 |
| Covert 3-CRT | 1.65 ± 0.18 | 460 ± 4 | 2.19 | 0.92 | 2.12 ± 0.23 | 456 ± 4 | 3.21 | 0.94 |
| Overt 3-CRT | 2.52 ± 0.24 | 443 ± 5 | 2.34 | 0.94 | 2.60 ± 0.26 | 448 ± 4 | 0.18 | 0.99 |

Experiment 1b (Letter) fit parameter data
|  | 0, -10, -20, -30, -40 (LHM) |  |  |  | 0, 10, 20, 30, 40 (RHM) |  |  |  |
| --- | --- | --- | --- | --- | --- | --- | --- | --- |
| Protocols | Slope (ms/°) | Intercept (ms) | $\chi^2_v$ | R | Slope (ms/°) | Intercept (ms) | $\chi^2_v$ | R |
| Covert SRT | 0.07 ± 0.07 | 203 ± 2 | 1.96 | 0.25 | -0.05 ± 0.07 | 207 ± 2 | 0.48 | 0.14 |
| Overt SRT | 0.36 ± 0.09 | 205 ± 2 | 1.42 | 0.62 | -0.06 ± 0.14 | 218 ± 2 | 1.02 | 0.67 |
| Covert 2-CRT | 4.57 ± 0.37 | 453 ± 5 | 0.14 | 0.99 | 4.26 ± 0.32 | 451 ± 5 | 0.63 | 0.98 |
| Overt 2-CRT | 3.79 ± 0.15 | 466 ± 3 | 0.46 | 0.99 | 3.91 ± 0.11 | 469 ± 3 | 0.99 | 0.99 |
| Covert 3-CRT | 5.91 ± 0.42 | 563 ± 7 | 0.76 | 0.98 | 5.62 ± 0.35 | 555 ± 6 | 0.19 | 0.99 |
| Overt 3-CRT | 4.37 ± 0.16 | 561 ± 4 | 1.02 | 0.98 | 3.90 ± 0.17 | 564 ± 4 | 0.58 | 0.98 |

Experiment 1c (Face) fit parameter data

**Table (2).** Correlation coefficient *R*^2^, reduced *χ*^2^ for the slope and intercept of regression linesin Figure 2(D, E, F). The data are aggregated over all participants (N= 13 for Experiment 2a, 2b; N = 12 for Experiment 2a eye tracking; N= 40 for Experiment 2c), categorized by protocol and subdivided into direction of scanning.

|  | 0, -15, -30, -45, -60 (LHM) |  |  |  | 0, 15, 30, 45, 60 (RHM) |  |  |  |
| --- | --- | --- | --- | --- | --- | --- | --- | --- |
| Protocols | Slope (ms/°) | Intercept (ms) | $\chi^2_v$ | R | Slope (ms/°) | Intercept (ms) | $\chi^2_v$ | R |
| Covert SRT | 0.45 ± 0.05 | 218 ± 2 | 1.89 | 0.88 | 0.27 ± 0.05 | 218 ± 2 | 2.07 | 0.72 |
| Overt SRT | 0.56 ± 0.05 | 249 ± 2 | 2.10 | 0.85 | 0.69 ± 0.06 | 242 ± 2 | 2.32 | 0.85 |
| Covert 3-CRT | 1.58 ± 0.09 | 399 ± 3 | 3.20 | 0.97 | 2.07 ± 0.10 | 383 ± 3 | 1.27 | 0.99 |
| Overt 3-CRT | 2.54 ± 0.14 | 392 ± 4 | 1.36 | 0.98 | 2.91 ± 0.14 | 386 ± 3 | 0.74 | 1.00 |

Experiment 2a (BBB-H) fit parameter data
|  | 0, -15, -30, -45, -60 (LHM) |  |  |  | 0, 15, 30, 45, 60 (RHM) |  |  |  |
| --- | --- | --- | --- | --- | --- | --- | --- | --- |
| Protocols | Slope (ms/°) | Intercept (ms) | $\chi^2_v$ | R | Slope (ms/°) | Intercept (ms) | $\chi^2_v$ | R |
| Overt 3 CR SET | 2.37 ± 0.10 | 135 ± 4 | 0.02 | 1.00 | 2.34 ± 0.07 | 142 ± 3 | 1.81 | 0.99 |
| Overt 3 CR SST | 0.60 ± 0.15 | 119 ± 5 | 0.37 | 0.92 | 0.55 ± 0.08 | 128 ± 3 | 1.31 | 0.91 |
| Overt SR SET | 1.57 ± 0.10 | 165 ± 4 | 1.40 | 0.97 | 1.98 ± 0.09 | 151 ± 4 | 0.07 | 1.00 |
| Overt SR SST | 0.26 ± 0.11 | 147 ± 4 | 1.16 | 0.60 | 0.42 ± 0.08 | 142 ± 3 | 0.76 | 0.89 |

Experiment 2a (Eye tracking) fit parameter data
|  | 0, -10, -20, -30, -40 (LVM) |  |  |  | 0, 10, 20, 30, 40 (RVM) |  |  |  |
| --- | --- | --- | --- | --- | --- | --- | --- | --- |
| Protocols | Slope (ms/°) | Intercept (ms) | $\chi^2_v$ | R | Slope (ms/°) | Intercept (ms) | $\chi^2_v$ | R |
| Covert SRT | 0.53 ± 0.09 | 216 ± 2 | 0.15 | 0.99 | 1.16 ± 0.10 | 210 ± 2 | 1.91 | 0.96 |
| Overt SRT | 0.58 ± 0.05 | 253 ± 2 | 1.63 | 0.90 | 1.30 ± 0.11 | 248 ± 2 | 0.98 | 0.97 |
| Covert 3-CRT | 2.89 ± 0.15 | 396 ± 3 | 0.97 | 0.99 | 3.55 ± 0.20 | 398 ± 3 | 1.82 | 0.97 |
| Overt 3-CRT | 3.13 ± 0.15 | 412 ± 3 | 3.90 | 0.98 | 5.09 ± 0.19 | 408 ± 3 | 2.92 | 0.99 |

Experiment 2b (BBB-V) fit parameter data
|  | 0, -10, -20, -30, -40 (LHM) |  |  |  | 0, 10, 20, 30, 40 (RHM) |  |  |  |
| --- | --- | --- | --- | --- | --- | --- | --- | --- |
| Protocols | Slope (ms/°) | Intercept (ms) | $\chi^2_v$ | R | Slope (ms/°) | Intercept (ms) | $\chi^2_v$ | R |
| Covert SRT | 0.75 ± 0.09 | 232 ± 2 | 0.47 | 0.98 | 0.89 ± 0.08 | 233 ± 2 | 0.55 | 0.99 |
| Overt SRT | 0.68 ± 0.10 | 233 ± 2 | 0.40 | 0.97 | 0.85 ± 0.10 | 234 ± 2 | 0.46 | 0.98 |
| Covert 2-CRT | 5.97 ± 0.26 | 493 ± 4 | 1.14 | 0.99 | 6.28 ± 0.23 | 489 ± 4 | 0.87 | 1.00 |
| Overt 2-CRT | 4.48 ± 0.13 | 497 ± 3 | 4.21 | 0.99 | 4.54 ± 0.15 | 497 ± 3 | 6.52 | 0.97 |
| Covert 3-CRT | 6.01 ± 0.31 | 623 ± 6 | 0.29 | 1.00 | 6.92 ± 0.33 | 625 ± 6 | 0.13 | 1.00 |
| Overt 3-CRT | 4.90 ± 0.19 | 613 ± 5 | 1.61 | 0.99 | 5.06 ± 0.18 | 610 ± 4 | 2.48 | 0.99 |

Experiment 2c (Face(b)) fit parameter data

**Table (3).** Correlation coefficient *R*^2^, and reduced *χ*^2^ for the slope and intercept of regression lines in Figure 1(G, H, I). The data are aggregated over all participants (N= 15), categorized by protocol and direction of scanning.

|  | 0, -15, -30, -45 (LHM) |  |  |  | 0, 15, 30, 45 (RHM) |  |  |  |
| --- | --- | --- | --- | --- | --- | --- | --- | --- |
| Protocols | Slope (ms/°) | Intercept (ms) | $\chi^2_v$ | R | Slope (ms/°) | Intercept (ms) | $\chi^2_v$ | R |
| Covert SRT | 0.03 ± 0.06 | 212 ± 2 | 0.99 | 0.17 | 0.12 ± 0.06 | 206 ± 1 | 6.12 | 0.26 |
| Overt SRT | 0.11 ± 0.06 | 213 ± 2 | 0.79 | 0.43 | 0.11 ± 0.08 | 214 ± 2 | 0.75 | 0.54 |
| Covert 2-CRT | 0.73 ± 0.09 | 337 ± 2 | 0.20 | 0.99 | 0.86 ± 0.07 | 330 ± 2 | 0.71 | 0.98 |
| Overt 2-CRT | 0.89 ± 0.08 | 337 ± 2 | 0.69 | 0.99 | 0.68 ± 0.06 | 340 ± 2 | 0.11 | 1.00 |
| Covert 3-CRT | 1.83 ± 0.13 | 370 ± 3 | 0.64 | 1.00 | 1.87 ± 0.11 | 373 ± 3 | 0.26 | 1.00 |
| Overt 3-CRT | 1.74 ± 0.12 | 377 ± 3 | 0.11 | 1.00 | 1.61 ± 0.12 | 377 ± 3 | 0.09 | 1.00 |

Experiment 3a (BBB) fit parameter data
|  | 0, -15, -30, -45 (LHM) |  |  |  | 0, 15, 30, 45 (RHM) |  |  |  |
| --- | --- | --- | --- | --- | --- | --- | --- | --- |
| Protocols | Slope (ms/°) | Intercept (ms) | $\chi^2_v$ | R | Slope (ms/°) | Intercept (ms) | $\chi^2_v$ | R |
| Covert SRT | 0.03 ± 0.06 | 212 ± 2 | 0.99 | 0.17 | 0.12 ± 0.06 | 206 ± 1 | 6.12 | 0.26 |
| Overt SRT | 0.11 ± 0.06 | 213 ± 2 | 0.79 | 0.43 | 0.11 ± 0.08 | 214 ± 2 | 0.75 | 0.54 |
| Covert 2-CRT | 0.66 ± 0.08 | 316 ± 2 | 2.73 | 0.91 | 0.47 ± 0.07 | 322 ± 2 | 1.25 | 0.90 |
| Overt 2-CRT | 0.73 ± 0.08 | 318 ± 2 | 0.71 | 0.98 | 0.67 ± 0.08 | 317 ± 2 | 0.33 | 0.99 |
| Covert 3-CRT | 1.09 ± 0.19 | 399 ± 3 | 1.20 | 0.96 | 0.98 ± 0.11 | 398 ± 2 | 0.04 | 1.00 |
| Overt 3-CRT | 1.14 ± 0.08 | 406 ± 2 | 2.62 | 0.96 | 0.68 ± 0.08 | 417 ± 3 | 0.29 | 0.99 |

Experiment 3b (Color) fit parameter data
|  | 0, -15, -30, -45 (LHM) |  |  |  | 0, 15, 30, 45 (RHM) |  |  |  |
| --- | --- | --- | --- | --- | --- | --- | --- | --- |
| Protocols | Slope (ms/°) | Intercept (ms) | $\chi^2_v$ | R | Slope (ms/°) | Intercept (ms) | $\chi^2_v$ | R |
| Covert CRT | 1.75 ± 0.11 | 287 ± 3 | 1.81 | 0.97 | 1.91 ± 0.11 | 291 ± 3 | 3.03 | 0.99 |
| Overt CRT | 1.94 ± 0.11 | 290 ± 3 | 2.35 | 0.99 | 1.91 ± 0.09 | 289 ± 2 | 1.83 | 0.99 |

Experiment 3c (LR) fit parameter data

### 2.2 Saccadic Eye movement for Overt Protocol

Single eye motion captured using a high-resolution camera is used to analyze the eye trajectories of each participant during the overt attention protocol in Experiment 3a. The saccade movement from the center of gaze to the peripheral stimulus for each trial is analyzed and plotted against time, where t=0 is when the stimulus appears. The raw data for a typical participant are plotted in Figure 5(A,C) for the SRT and CRT protocols. The trajectories are sorted and shifted to the average saccade start time for each eccentricity to produce the trimmed eye trajectory plots in Figure 5(B, D). The average saccade start and saccade end times were extracted from the eye trajectories for N=12 subjects and plotted in Figure 6. Data for one subject was discarded due to technical issues in data recording.

**Figure (5).**
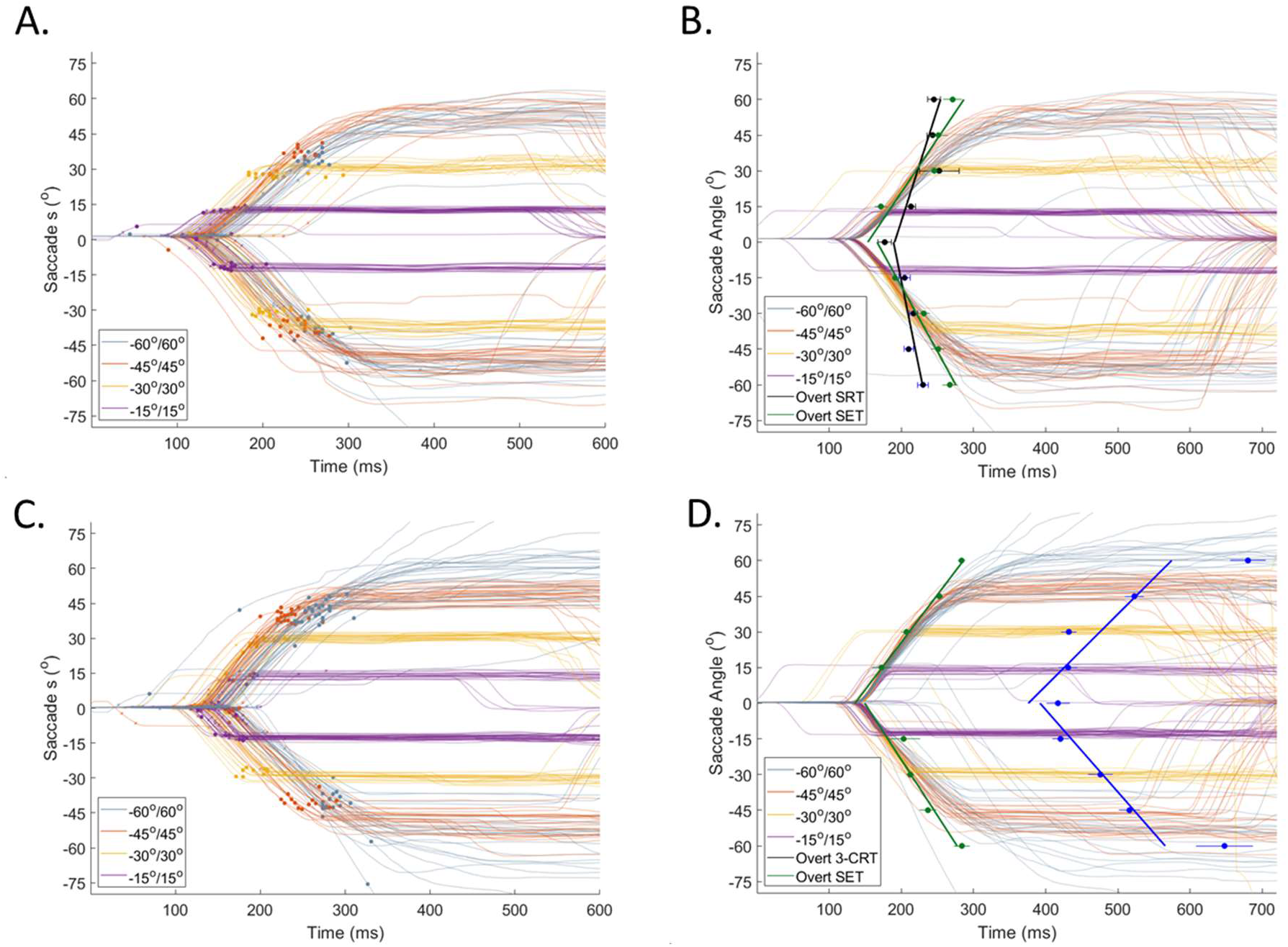
The centroid movement of the left pupil for a typical participant is plotted with respect to time for experiment1 SRT protocol (A, B) and 3-CRT protocol (C, D). (A, C) Raw eye trajectory for all trial with saccade end points. (B, D) Trimmed eye trajectory for all trial overlaid with mean RT and mean saccade end vs eccentricity data.

**Figure (6).**
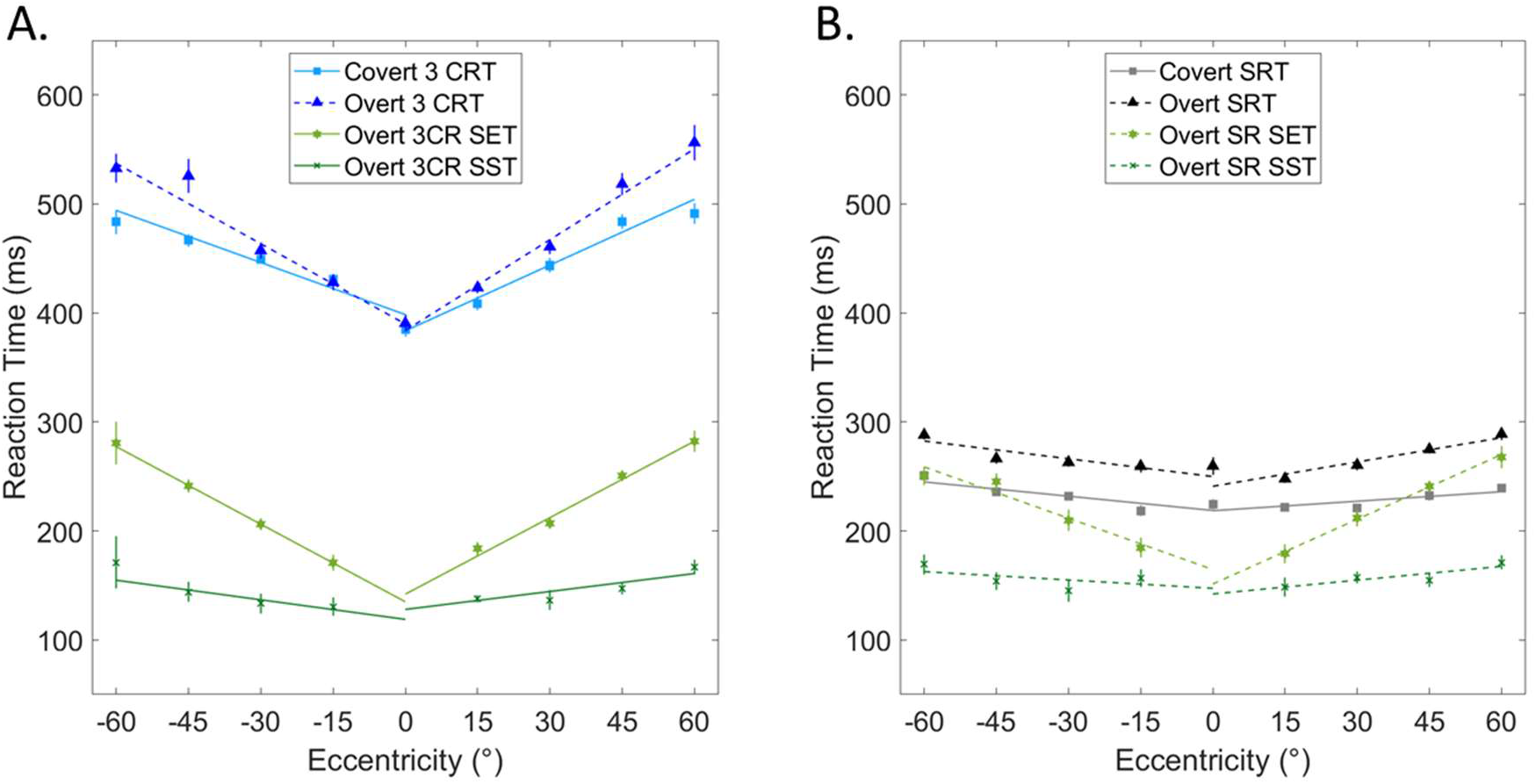
Mean reaction time, saccade end time and saccade start time vs eccentricity for overt protocol and reaction time for covert protocol for experiment 3c. The data are aggregated over all participants (N= 12). (A) Data shown for 3-CRT protocol. (B) Data shown for SRT protocol.

## 3 Discussion

### 3.1 Reaction time dependence on eccentricity

The experimental results verified our postulation that semantic spatial information is converted to temporal information. From the graphs in Figure 1 it is observed that when the participant(s) responded to only one target stimuli at different eccentricities, in both the overt and covert attention cases, RT was fairly independent of the eccentricity at which the stimuli appeared. The slight increase in reaction time with increasing eccentricity maybe attributed to the relative reduction of cone density in the periphery of the retina (Osterberg, 1935). This observation was expected since the reaction time was purely reflexive as the participant was not required to make a choice. When semantic information was introduced by presenting different stimuli and the participant was asked to respond accordingly, the overall RT increased. Additionally, we see that at larger eccentricities, reaction time increases linearly with eccentricity for the choice reaction experiments. We observed a significant correlation, with similar results for both covert and overt attention. Obviously, the gradient of the slope is participant dependent since several factors affect reaction time, but the correlation is significant across all participants. From the aggregated data presented in Figure 1 we observed similar trends between reaction time dependence for SRT and CRT protocol for all experiments. This result supports our hypothesis that a linear frame shift is required to remap our stored memory of objects to their eccentric locations in space. We hypothesize that this process utilizes alpha brainwaves, which are responsible for the 100-200 ms response delay between a stimulus appearing in the center of the visual field and at the periphery. Additionally, studying saccadic eye movements using the Overt attention protocol, we see that the mean gradient of the trajectories mimics that of the reaction times using the overt attention protocol, which in turn fairly overlaps with the covert attention protocol where there was no eye movement.

### 3.2 Reaction time for Overt vs Covert Attention

Analyzing the aggregated data for all participants, we observe that reaction time is generally the same for overt attention and covert attention for both choice and simple reaction times, except in the 3-CRT protocol for Experiment1c where covert attention produced a higher RT than did overt attention, and for Experiments 3a and 3b where overt attention produced a higher RT than did covert attention. For the central stimulus, there is generally no significant difference between RT for covert and overt attention, except in Experiments 3a and 3b where overt RT was significantly higher than covert RT for the SRT protocol. Therefore, the time required for saccadic eye movements that are employed to focus on a sudden stimulus onset are shown to be equal to the time delay created from the allocentric frame shift. In terms of daily life, this equivalence is consistent with how we view external space. When viewing an object with our peripheral vision, we obtain a perception of that object’s distance. Moving our head or eyes to recenter that object on our fovea must create the same perception of distance. Otherwise, this movement would cause our vision to be constantly shaky and unstable.

### 3.3 Reaction Time dependence on stimulus direction

From the aggregated data we observe good symmetry in change in choice RT with increasing eccentricity along both the right horizontal meridian and the left horizontal meridian for both overt and covert protocols across all experiments. Along the vertical direction, RT was faster in the lower vertical meridian than in the upper vertical meridian for overt attention and fairly symmetric for covert attention. This may be attributed to better visual performance in the lower vertical meridian (Liu et al., 2006).

### 3.4 Reaction Time dependence on stimulus complexity

From Experiments 1 and 3, we observe that RT increased when the number of possible semantic stimuli that could be presented increased, i.e. RT is higher for the 3-CRT than the 2-CRT protocols. Moreover, In Experiment 1, we observe that as complexity of target stimuli increases from gabor patterns to alphabetical letters, to unfamiliar faces, the slope of the RT-eccentricity relationship also increased. More complex stimuli appear to require more time to be retrieved from a subject’s memory and for comparison processes that feed into the final response decision.

### 3.5 Alternative Accounts of the relationship between RT and eccentricity

We are not the first to study the effects of eccentricity of a visual onset on RT of a behavioral response. There is a long history of empirical work dating back to more than a century. Nevertheless, the primary conclusion of this historical body of work is that RT differences are due largely to changes in density of retinal cells across the surface of the retina. Cones are packed more tightly in the fovea while rods are more sparsely distributed in the non-foveal parts of the retina. Thus, there are more light-sensitive photoreceptors in the fovea and fewer at an increasing distance from the fovea. The decrease in density of photoreceptors is the presumed explanation for the effects of eccentricity of the visual stimulus to reaction time, with slower RTs at regions of the retina that have fewer photoreceptors. While this account has some explanatory power, it fails to account for one of our major findings, almost no effect on RT of eccentricity in the SRT procedures, but large effects of eccentricity on RT in the CRT procedures. Presumably, different densities of photoreceptors would have the same effect on SRT and CRT procedures, which was not the case. Only our theory of SRT being solved by unconscious processing of a sudden stimulus onset not requiring the time-code dependent NHT on the one hand, but CRT being solved by conscious processing requiring the time-code dependent NHT process on the other, can explain the difference in the RT-eccentricity relations found between these two protocols.

## 4 Methods and Materials

### 4.1 Participants

Due to the Covid-19 pandemic, no external participants were recruited for the study. Due to these circumstances,all participants acted as both subjects and experimenters and were therefore not blind to the hypotheses or methods. For Experiment 1, data were instead collected from internal members (7 females, 6 males, aged 17-22) of the laboratory, and subjects conducted the experiments remotely using a standardized setup at their residential homes. Experiment 2 was similarly conducted remotely with 15 participants (8 females, 6 males, aged 18-22). For Experiment 3, data were collected from 13 participants (6 female, 7 males, aged 18-32) affiliated with the laboratory, 12 undergraduate researchers and 1 post-doctoral researcher. Data were collected from each participant in 2 separate sessions at UCLA campus. Experiment 3a was conducted in the first session and experiment 3b was conducted in the second session. The participants were all healthy, right-handed individuals with normal or corrected-to-normal vision.

Participants were drawn from the undergraduate and graduate UCLA student population in accordance with approved procedures from the Institutional Review Board (IRB # 19-001472). The experiment was initially developed on campus in early 2020. Due to the COVID-19 pandemic, data-taking was conducted remotely with internal members from the summer of 2020 until the summer of 2021. Afterwards, standardized experimental setups were developed in-lab and external participants were recruited for additional, professional data-taking. These groups’ data were assessed separately for consistency, then combined in an aggregate analysis as shown in the Results.

### 4.2 Setup- TV based

The primary and ideal setup for the experiment consisted of a 55” or greater TV connected to the subject’s computer via HDMI cable. The subject then sat with a 50 cm distance from the subject’s eyes to theTV’s center marked by a white cross mark cue, using a chin rest to position the subject’s head. The chin rest stabilized head movement during the experiment and maintained the subject’s eye level at the middle of the display.A keyboard was used for experimental input. Subject reaction time was recorded after a subject pressed the designated key corresponding to the stimulus that appeared on the screen. All subjects took data in a well-lit room.

Subjects that were not able to take data under the primary setup utilized an alternative setup. This consisted of using an external computer monitor (connected via HDMI) or the laptop screen itself. However, for subjects with a smaller screen size (less than 55”) to display stimuli at the full range of eccentricities that were used in the experiments, subjects had to decrease their distance to the screen. Additionally, these subjects took data in two separate sessions, one with negative eccentricities and the other with positive eccentricities. Instead of stimuli appearing on either side of the center cue as they would for a large TV, the “central” cue would be located on the rightmost side of the display with the stimuli appearing to the left to test negative eccentricities. When testing positive eccentricities, the cue would be located on the left with stimuli appearing to the right.These subjects were positioned directly in front of the cue on the left or right side of the screen depending on which eccentricities were being tested. This allowed subjects with an insufficiently large screen to record data that matched data recorded by subjects using a large TV. All subjects performed the experiments on screens that exceeded a 60 Hz refresh rate to minimize errors associated with timing. The typical set-up is shown in Figure 7.

**Figure (7).**
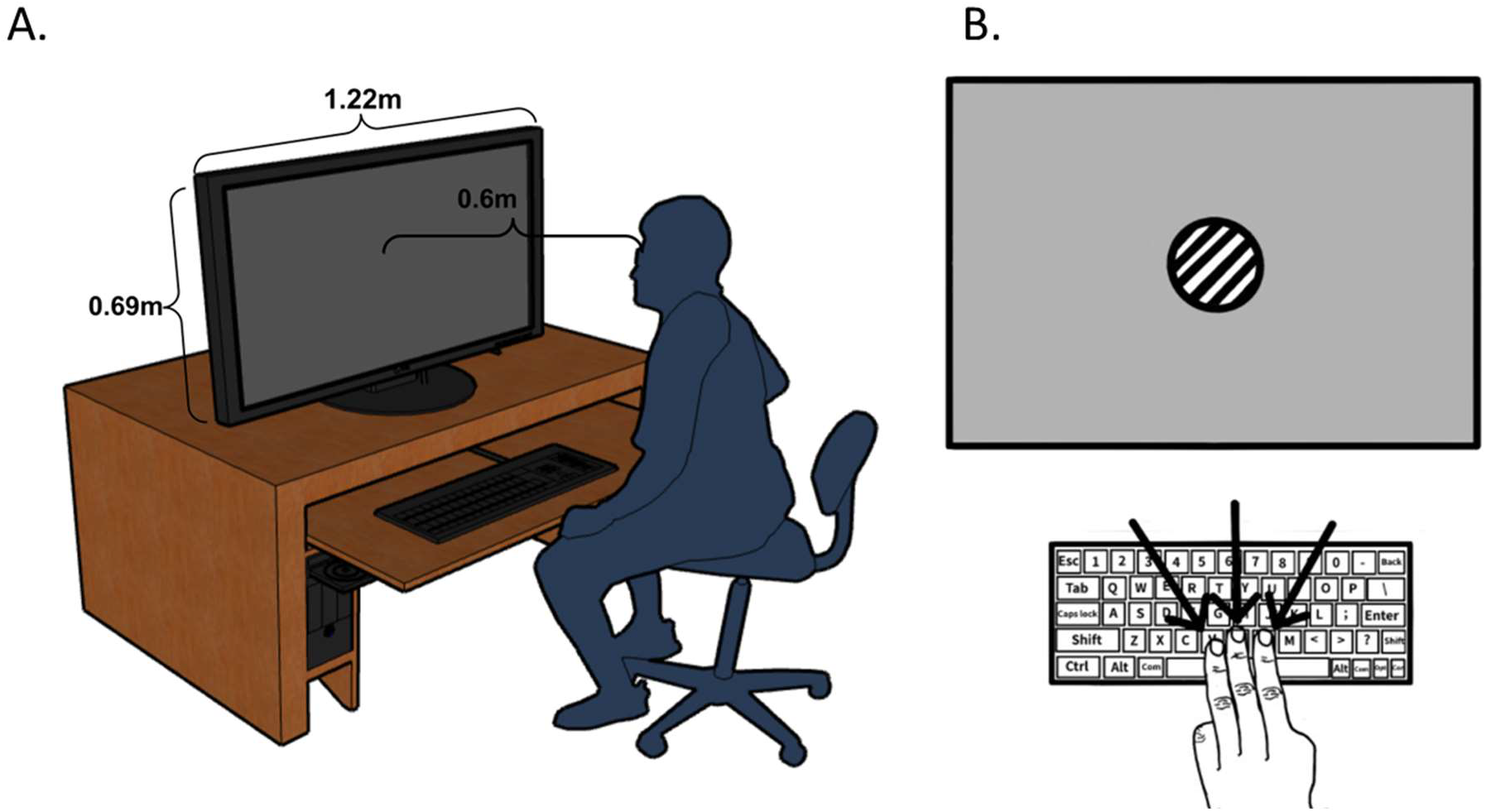
A typical setup for subjects in Experiment 1. A chin rest (not shown) was used to steady subject head movement. A keyboard was used to record subject input. Subjects sat 50cm away from the center of the laptop screen to view the various visual stimuli during the test sessions. (A) 55” TV setup. (1 (B) Diagram of screen with keyboard for subject input.

### 4.3 TV-based experiment stimuli set

The description of the stimuli sets are given below.

Gabor-A circular gabor pattern was shown at varying eccentricities and varying angles or rotation. The stimulus was generated using the GratingStim component of PsychoPy. The stimuli had a gaussian mask, a size of four degrees, a spatial frequency of 1.25/degree, and a contrast of one. For the experiments where one stimulus was shown, the gabor pattern was shown with the grating displayed vertically. For experiments where there were two stimuli, the gabor pattern could appear vertically or horizontally (or rotated 90 degrees). For three stimuli, the patterns were displayed vertically, horizontally, or at a 45 degree angle.

Alphabetical Letters (EPB)-A familiar letter was displayed at varying eccentricities. Three different letters were utilized in this experiment: “E”, “P”, and “B”. For the experiments where one stimulus was displayed,the letter “E” was shown. For the experiments that had two stimuli displayed, “E” and “P” were shown. For the experiments that had three stimuli displayed, “E”, “P”, and “B” were shown. This experiment was performed with covert and overt conditions (same conditions as in the face experiment were followed).

Unfamiliar Face-An unfamiliar face image was displayed at varying eccentricities. The various face images utilized in this experiment were selected from a collection of face images (DeBruine & Jones, 2017). These images were removed of color and were resized to the same size. For the experiments where one stimulus was displayed, a singular face was shown. For the experiments in which the stimulus that was shown on each trial was drawn from a set of three, the face from the single stimulus experiment was shown along with an additional face image. For the experiments that had three stimuli displayed, the faces from the two stimuli experiments were shown along with an additional face image. All the face images utilized in this experiment were distinct and different from one another.

### 4.4 Setup- LED based

Experiment 3 was conducted in a large 7m by 8m room in the UCLA Physics and Astronomy department. The room had ambient lighting provided by in-ceiling florescent lights, incandescent lights throughout the duration of the experiment. The experimental setup used an upright PVC geodesic hemisphere with six 5m long individually addressable LED Strips (ALITOVE Electronic Technology Co, Shenzhen, Guangdong, China) assembled 30 degrees apart in radial arrangement. A 3D rendering of the setup is shown in Figure 8. The setup was at the end of the room with a computer setup next to it for the experimenter to conduct the behavioral procedures. An Arduino Mega 2560 integrated circuit(Arduino LLC, Boston, Massachusetts, USA) was programmed to control each LED strip. The horizontally aligned strip was used for Experiment 2a and the vertically aligned strip was used for Experiment 2b. A chin and forehead rest was placed 1 meter off the ground and 1 meter away from the LED strips, affixed to the inside surface of thePVC hemisphere. Participants were seated in a chair which adjusted their height to match the level of a fixed chin and forehead rest. The forehead rest was further adjusted to minimize head movement, and the chin rest was also adjusted to level participants’ eyes with the horizontal meridian of the dome.

**Figure (8).**
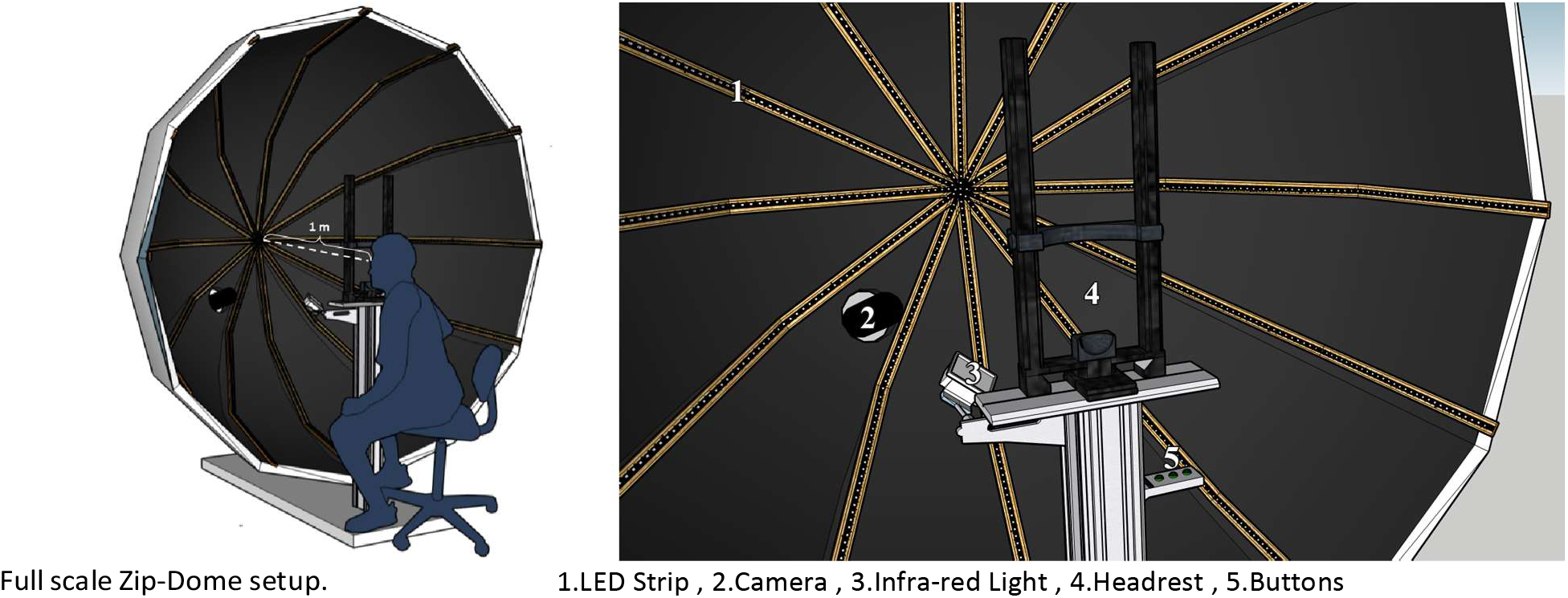
Zip-dome setup with attached LED-strips. The hemisphere shape of the dome ensures that all spatial locations at which the stimulus is provided is at a 1 m distance from the participant’s eye. The participant is instructed to press the first button if they observed one LED, press the second button if they observed two LEDs and press the third button if they observed three LEDs using their index, middle and ring finger respectively. The camera tracks the saccadic movement of the eye illuminated by the IR light.

The LED strips used in the experiment were WS8212B individually addressable LED strips with 200 LEDs within each strip. The adjacent LEDs were 1.5cm apart with a sector angle of 0.833 degrees. Three Arduino buttons were secured to a breadboard which was wired to the Arduino controlling the experiment. The buttonpress was recorded by the Arduino to measure the reaction time in the Arduino Serial Monitor integrated with LabView, which was used in calculating reaction time.

Single eye motion is captured using Mako U-029B sCMOS sensor (Allied Vision, Exton, Pennsylvania, USA) fitted with a variable zoom lens to adjust for the field of view. The sensor was placed 40^°^ below the LHM and 25 cm from the center of the apex of the dome. The sensor was used to ensure that all participants had their eye at the center and to track the saccadic movement of the left eye. Sitting at the opposite position of the participant,the camera has a near head on view of the participant’s eye motion. The camera operates at 245 frames per second with an exposure time of 3400 *μ*s, capturing the reflected infrared light by the illumination light source.Synchronization between the camera and the experimental Arduino Mega microcontroller is performed by a master Arduino Uno Rev. 3 board which awaits a signal from the experimental Arduino Mega at a clock rate of 490 Hz while also triggering the camera via a Transistor-transistor Logic (TTL) signal. The separate components of the setup are detailed in Figure 8.

### 4.5 Behavioral Task

#### General Procedure- TV based

Prior to data collection, subjects calibrated their screen using a Python script. This calibration recorded all necessary information, including screen width (pixels and centimeters), screen height (pixels), left and right screen edges, screen center, and multipliers for letter height, circle radius, horizontal stimulus location (left and right eccentricities), and image display dimensions. All protocols used this calibration to accurately determine the size and location of the stimuli. All protocols and this calibration were written in Python3 and were displayed using PsychoPy (Peirce et al., 2019). All protocols and the calibration will be made freely accessible.

To account for system latency, we set up ten trials on PsychoPy that would flash a Gabor pattern and wait for a response (hitting the “V” key). The subjects had to record these trials with a slow-motion camera (240 Hz) and determine their reaction time for each of the ten trials through the video. They subtracted these times for each trial from the respective PsychoPy times (PsychoPy RT - Video RT). The average difference is the latency (to be subtracted from all future PsychoPy RTs to get the adjusted value).

The eccentricity experiments were designed to show that determination of semantic shapes had eccentricity dependence. To do this, we used varying semantic information at different eccentricities and recorded the reaction times of the subjects’ responses. We started off with one stimulus shown at varying eccentricities to test simple reflexive responses, which we expected to be the same regardless of stimuli.

All stimuli were shown at locations ranging from zero degrees to forty degrees left or right from the center,in increments of five or ten degrees. For the Gabor and Unfamiliar Face experiments, stimuli were presented at 0, 5, 10, 15, 20, 25, 30, 35, and 40 degrees from the center; for the Familiar Character experiments,stimuli were presented at 0, 10, 20, 30, and 40 degrees away from the center. The reason for the change in stimulus locations was to prevent the participants from becoming fatigued. All locations were able to be shown on a large TV keeping the center at the exact center of the screen. Subjects with a smaller screen chose the center to be on the left or right, allowing the full range of eccentricities to be shown as described in Procedural Setup. The size and locations of the stimuli were calculated in accordance to the individual subject’s distance to the screen.

There were a total of ten trials taken at each eccentricity location with twenty taken at the center, resulting in a total of 180 trials when in five degree increments and 100 trials when in ten degree increments. In all experiments, a white cross hair pattern would appear first for 100 ms. This white cross ensured that subjects were looking at the center of the screen at the start of each trial. This was followed by an interstimulus interval (ISI) that was random, ranging from 300 ms to 800 ms. This ensured that subjects were not able to predict when the stimulus would appear. The ISI was followed by the presentation of the stimulus, which would remain on the screen until a response from the subject. Time until response was recorded and saved as data if the response was correct, and if the reaction time was not greater than 1500 ms. Any incorrect responses or responses that exceed the designated reaction interval would not be counted, and information about the stimulus and its location would be saved to be tested again following completion of the first set of trials. After going through all the trials that were programmed, the subject then retakes data from the mistakes until no more errors are made. This allowed all subjects to contribute the same amount of data for each eccentricity condition despite incongruences in the number of mistakes made.

In the eccentricity experiments, the same key mappings were used throughout, with “V” used for experiments with one stimulus shown, “V” and “B” used for experiments with two stimuli shown, and “V”, “B”, and “N” used for experiments with three stimuli shown. This allowed for all keys to be pressed with one hand, which the subject could choose whether to use their left or right hand based on comfort or handedness.

For experiments that presented a higher degree of difficulty, an audible beep was implemented and sounded whenever the subject’s response to the displayed stimulus was incorrect. The audible beep was used as an indication of inaccuracy to help allow the subject to distinguish and differentiate one stimulus from another.This beep was used in our “Unfamiliar” experiments, including Unfamiliar Faces and Unfamiliar Characters.

#### General Procedure- LED based

For BBB experiments (2a, 2b, 3a), each session consisted of the presentation of more than 200 trials of either one blue flashing LED, two adjacent blue flashing LEDs or three adjacent blue flashing LEDs at different locations along the horizontal or vertical LED strip, shown in Figure 8. For the color experiment (3b), the stimulus presented was either a red LED, blue LED, or yellow LED. For the LR experiment (3c), the semantic information was the varying order in which two LEDs flashed. Subjects were instructed to press buttons corresponding to the order in which two adjacent LEDs flashed, separated by 150 ms, as quickly as possible.

Each reaction time trial started with a 200ms center white cross hair stimulus to prompt the participants to align their gaze to the center of the LED strip. For Experiment 2, the pre-trial cue was 3 adjacent white LEDs at the center. The central cue was also used to prime the participant to optimize reaction time and reduce error in response (Bertelson, 2018). The Arduino controlling the experiment selected a random time period between 300 and 800 ms for which no stimulus was present, followed by the target stimulus.This variable interstimulus interval (ISI) was chosen to minimize the likelihood of having participants’ photo receptors in a refractory state at target stimulus onset and to minimize the predictability of target stimulus onset which could bias participant response. At the end of this time period, the randomly chosen stimulus turned on at a randomly chosen eccentricity from a preset list via the Arduino random number generator. The list of possible angles were specific to the experiment. Each eccentricity was tested an approximately equal number of times with a maximum difference of 10 trials. The mean number of trials per eccentricity was 22. In Experiment 2(a, b, c), the stimulus was randomly presented at 0, 15, 30, or 45 degrees eccentricities, as measured from the center fixation point, along either the left or right horizontal meridian. In Experiment 3a, the stimulus was randomly presented at 0, 15, 30, 45, or 60 degrees eccentricities, as measured from the center fixation point, along either the left or right horizontal meridian. The adjacent blue flashing LEDs were approximately 4 LEDs or 6 cm apart for both 3 and 2 adjacent flashing LEDs. In Experiment 3b, the stimulus was randomly presented at either 0, 10, 20, 30, or 40 degrees, as measured from the center fixation point, along the lower or upper vertical meridian. A smaller range of eccentricities were tested in Experiment 2 since the human binocular visual field is limited to 50 degrees above the meridian and 70 degrees below the meridian. The adjacent blue flashing LEDs were approximately 3 LEDs or 4.5 cm apart from each other for 3 adjacent flashing LEDs and 4 LEDs or 6 cm apart for 2 adjacent flashing LEDs. The gap between adjacent LEDs was less than that in Experiment 1, since the eccentricities at which the stimulus was presented were spaced more closely together. The target stimulus remained on until one of three criteria was met: 1) The participant pushed the button correctly corresponding to the given stimulus, 2) the participant pushed the button not corresponding to the given stimulus (i.e., an error of commission), or 3) the participant did not give a response within 1 second after the stimulus onset (i.e., an error of omission). Examples of 2 subsequent trials for Experiments 2a, 3a, and 3b are shown in Figure 9 and examples for Experiments 3b and 3c are shown in Figure 10. The participant was not given any feedback with regards to accurate or inaccurate responses because such feedback has been shown to slow down processing of the subsequent target stimulus (Koehn et al., 2008). Trials with errors were excluded from data analysis. The push buttons were located on the participant’s right-hand side fixed to the structure supporting the chin and forehead rest (see Figure 8). The participant’s response was recorded by the Arduino Serial Monitor and stored in LabView (National Instruments,Austin, Texas, USA). Upon meeting one of these three criteria, the target stimulus would turn off. No stimulus was presented for a minimum inter-trial interval (ITI) of 500 ms, from the termination of the target stimulus onthe antecedent trial to the onset of the orienting stimulus on the subsequent trial. Each participant was given ascheduled 200 trials. An additional trial, randomly chosen from the set, was added for each incorrect response, or missed response to ensure sufficient statistics. At the conclusion of the study session, three blue LEDs flashed along the center of the strip in ascending order to notify the participant that the session was complete.

**Figure (9).**
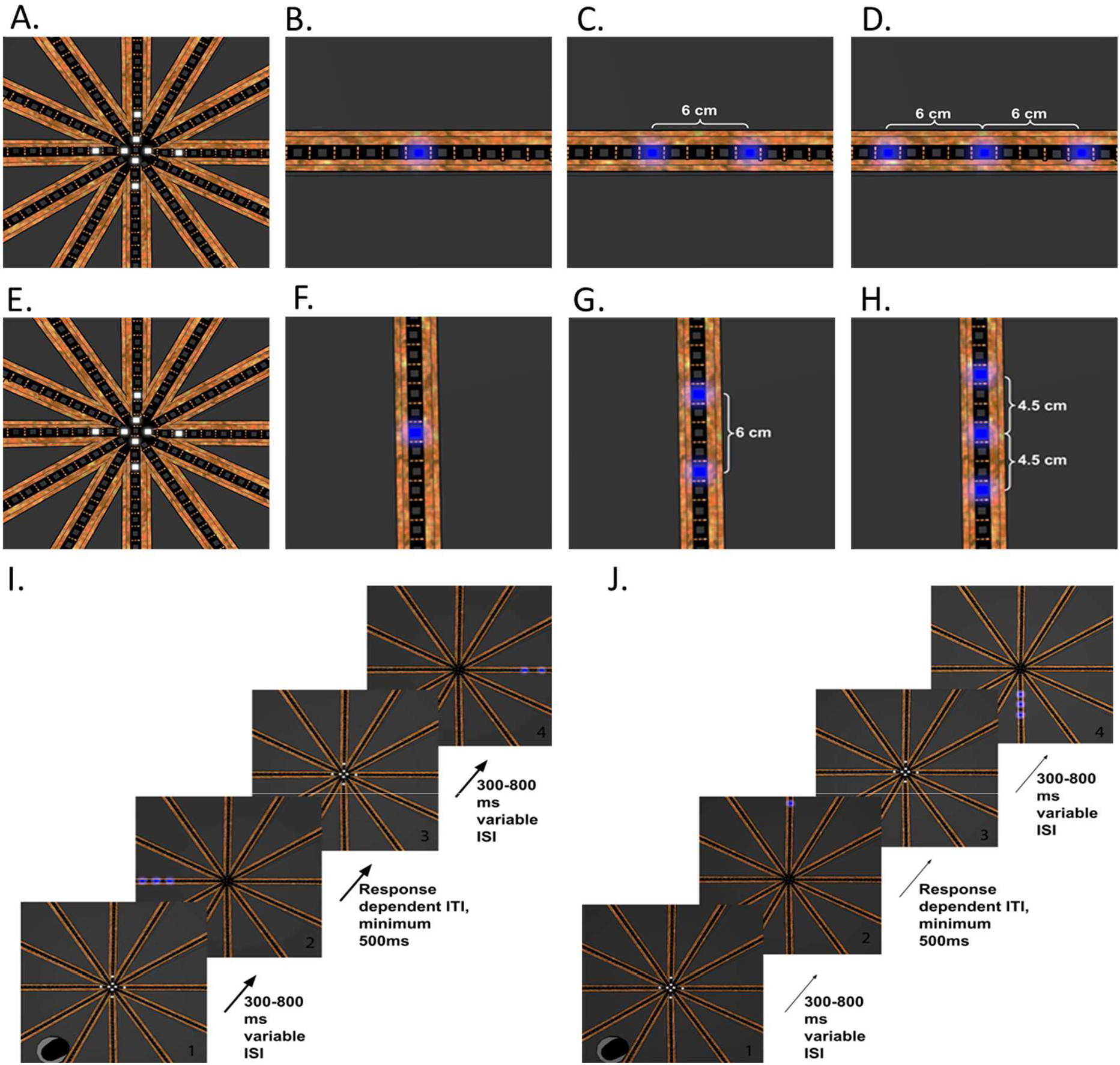
(A, E) White Cross Hair exogenous pre-trial cue. (B, F) Single LED target stimulus. (C, G) Three adjacent LEDs target stimulus. (B, C, D) Visual stimulus for Experiments 1 and 3. The random target stimuli would appear at a randomlocation along the horizontal meridian. (F, G, H) Visual stimulus for Experiment 2. The target stimulus is centered about the defined eccentricity location. (I) Sequential trial example in Experiments 1 and 3. (J) Sequential trial example in Experiment 2. Each trial is initiated by white cross hair cue at the center (1, 3) for 200ms. After a variable ISI, a random target stimulus (2, 4) appears at a random location along the horizontal meridian in Experiments 2(a) and 3(a) or along the vertical meridianin Experiment 2(b). The duration of the target stimulus and ITI is dependent on the three criteria described in the general procedure section.

**Figure (10).**
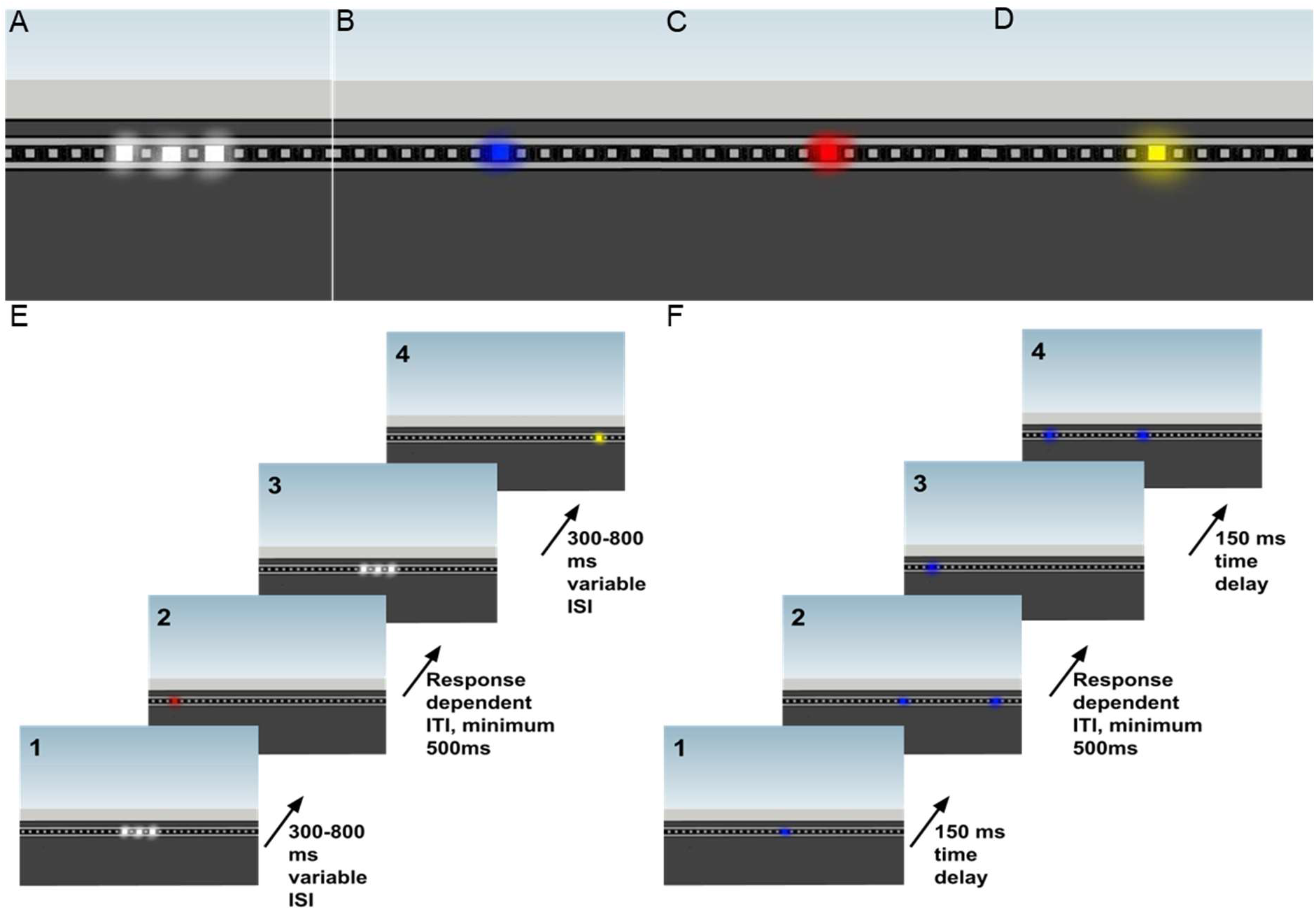
(A) White exogenous pre-trial cue. (B) Single blue LED target stimulus. (C) Single red LED target stimulus. (D) Single yellow LED target stimulus. The random target stimuli would appear at a random location along the horizontal meridian. The target stimulus is centered about the defined eccentricity location. (E) Sequential trial example in Experiment 3b. (F) Sequential trial example in Experiment 3c. In Experiment 3b each trial is initiated by white cue at the center (1, 3) for 200ms. After a variable ISI, a random target stimulus (2, 4) appears at a random location along the horizontal meridian. In Experiment 3c, a blue LED appeared randomly either at the center or at some designated eccentricity at the periphery followed by a second blue LED at the other location after a 150 ms delay. The duration of the target stimulus and ITI is dependent on the three criteria described in the general procedure section.

#### Eye Tracking Methods

For Experiment 2(a, b) the experimenter was seated out of view adjacent to the dome apparatus and monitored the eye movement via real time imaging from the eye tracking camera. For Experiment 3b, the single eye movement of participants was monitored by the experimenter but not recorded for saccadic movement analysis.Once the target stimulus appeared, the master control board received the signal that the experiment board had begun the experimental trial and relayed a serial communication signal to the camera control software to begin trial video capture. The camera recorded for 2500ms after the stimulus onset or for 1000ms if the participant did not press a button. The camera control software was custom made in LabVIEW and appended each additional trial onto a multipage tiff (MTIFF) file with additional trial metadata for later access. Upon completion, the MTIFF file was then broken up into the individual trials for analysis.

#### Experimental Protocols

For Experiment 1(a, b, c) and Experiment 3(a, b), there were a total of six different experimental conditions with three conditions testing covert attention and three testing overt attention in Experiments 1 and 2 and four experimental conditions in Experiment 3. For the overt condition, the participant was allowed to move their eyes freely to the location of the stimuli and refocus on the white-flashing cross stimulus before the next trial began. For the covert attention condition, participants were asked to keep their eyes focused on the center of the setup for the duration of the experiment. In Experiment 3, the experimenter monitored eye movement via real time imaging from the eye tracking camera. The experiment was further subdivided into three different tasks for each attention condition, one testing for Simple Reaction Time (SRT) and the other two testing Choice Reaction Time (CRT). The six protocols are summarized in Table 4. In protocols 1 and 2, a single target stimulus was randomly presented at various eccentricities and the subject was asked to press the first button using their index finger when they observed the target stimulus. In protocols 3 and 4, the target stimulus was randomly chosen from the first two in the set and in protocols 5 and 6, the target stimulus was randomly chosen for the full set of three.

**Table (4).** Summary table comparing stimulus and attention behavior across different protocols. Protocols 1 and 2 tested for SRT under covert and overt attention conditions, respectively. Protocols 3 and 5 tested for choice RT under the covert attention condition, and protocols 4 and 6 tested for choice RT under the overt attention condition. All other parameters, such as LED brightness, LED color, center cue length, and trial length were controlled across all protocols.

|  | Protocol Parameters |  |  |  |
| --- | --- | --- | --- | --- |
|  | Overt | Covert | SRT | CRT |
| Protocol 1 | x |  | x |  |
| Protocol 2 |  | x | x |  |
| Protocol 3 | x |  |  | x |
| Protocol 4 |  | x |  | x |
| Protocol 5 | x |  |  | x |
| Protocol 6 |  | x |  | x |

The target stimulus was randomly presented at various eccentricities and the subject was asked to press the firstbutton using their index finger when they observed the first stimulus type, the second button when they observed the second stimuli type and the third button when they observed the third stimulus type. The participants performed all six different tasks for each experiment on the same session in random to avoid training effects. For experiment 3(a, b), the subjects only performed protocols 1, 2, 5, and 6.

For experiment 2c, there were only two experimental conditions testing covert and overt attention. The subject pressed the left button if the relatively left LED flashed first and the right button if the relatively right LED flashed first.

### Statistical Analysis for RT data

Reaction time trial data were categorized by their eccentricities, and further analysis was conducted within the dataset separated by each eccentricity angle.

#### Outlier Analysis

Outlier data were removed before averaging the RT at each eccentricity. Data points that fell below the 25th percentile minus 1.5 times above the interquartile range or above the 75th percentile plus 1.5 times the interquartile range for reaction time data at each angle were removed. The resulting data points were then averaged and used in chi square calculations.

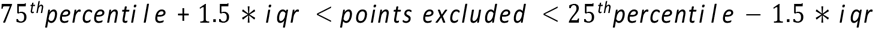

#### Error Analysis

The resulting reaction times were averaged, and error bars were plotted using the calculated standard error,taking the standard deviation of the eccentricity angle reaction time divided by the square root of the number of trials within the eccentricity angles. After reaction time data were categorized by angle and outliers were removed, the standard deviation and standard error were calculated, with the standard error being used in the plots.

#### Hypothesis Testing

To ensure that an attention shift occurred during each choice RT block, response accuracy was analyzed using aone sample t-test for a mean of ^1^, with 3 being the number of available choices for the 3-CRT protocol. The test was performed for each angle and subject’s data was removed for a given angle if the corresponding p-value was greater than 0.05.

#### Normalization of average RT

To better represent the relationship between RT and eccentricity in the aggregated analysis, we normalized each participant’s data to a global average for each protocol. We normalize the data by subtracting the appropriate participants mean performance from each observation, and then add the grand mean score to every observation. Let y be the i-th participants score in the j-th condition (i = 1, … N and j = 1, … M). Then define the normalized observations z-

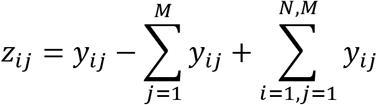

#### Line of best fit and correlation coefficient (*χ*2)

A best fit line was added using a least-squares linear regression for both positive and negative angles and their corresponding reaction times, with 0 degrees being included in both positive and negative fits. The linear regression returned slope and intercept parameters as well as the correlation coefficient and a two-sidedp-value in which the null hypothesis assumes a slope of 0. Additionally, the standard error of the fit was returned.

A best fit line of the form y = mx + b was calculated using chi square minimization to find the best fit slope and intercept parameters for both positive and negative angles with each side including 0 degrees. From the chi square minimization, error estimates for the slope and intercept parameters were computed by holding the other parameter constant at the best fit and calculating which parameter values would yield the minimum chi square value + 1. This process was done for both slope and intercept parameters. The reduced chi square was also calculated to determine goodness of fit by dividing the minimum chi square value by the degrees of freedom. The correlation coefficient was also calculated using the Numpy library in Python.

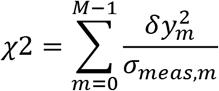

where 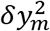 is defined as the mth residual 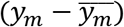 and *σ*_*meas,m*_ is the standard error of the reaction time data for an angle m.

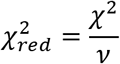

where *ν* is the degrees of freedom.

## 5 Acknowledgments

A total of ^~^70 extra students helped us as participants in various stages of RT experiments in 2020 – 2022. We thank them for their kind commitment.

This work was in part supported by the Dean’s office of life science, Dean’s office of physical science, Chair’s office of the department of physics and astronomy, and the Instructional Improvement grant by the Center for the Advancement of Teaching, all at the University of California, Los Angeles.

## 6 Contribution

KA developed the concept and supervised the project. UA, JC, CK, AD, DE, TM and several other undergraduate researchers designed and constructed the hardware. Patrick Wilson, TM and BT wrote the PsycoPy protocol. UA, JC, CK and TM wrote the Arduino protocol. DE managed the IRB. All listed authors coordinated the data taking. UA, JC, AD, DE, TM and BT analyzed the data. IB and UA designed the 3D setup models and experiment protocol diagrams. UA wrote the original manuscript. KA and AB edited and proofread.

## Supplementary Material

For supplementary material accompanying this paper, please visit https://github.com/uafifa/RTvsEccentricity

## Notes

https://github.com/uafifa/RTvsEccentricity

### Competing Interest Statement

The authors have declared no competing interest.

