## Supplemental Information for "Visual Perception of 3D Space and Shape in Time - Part I: 2D Space Perception by 2D Linear Translation"

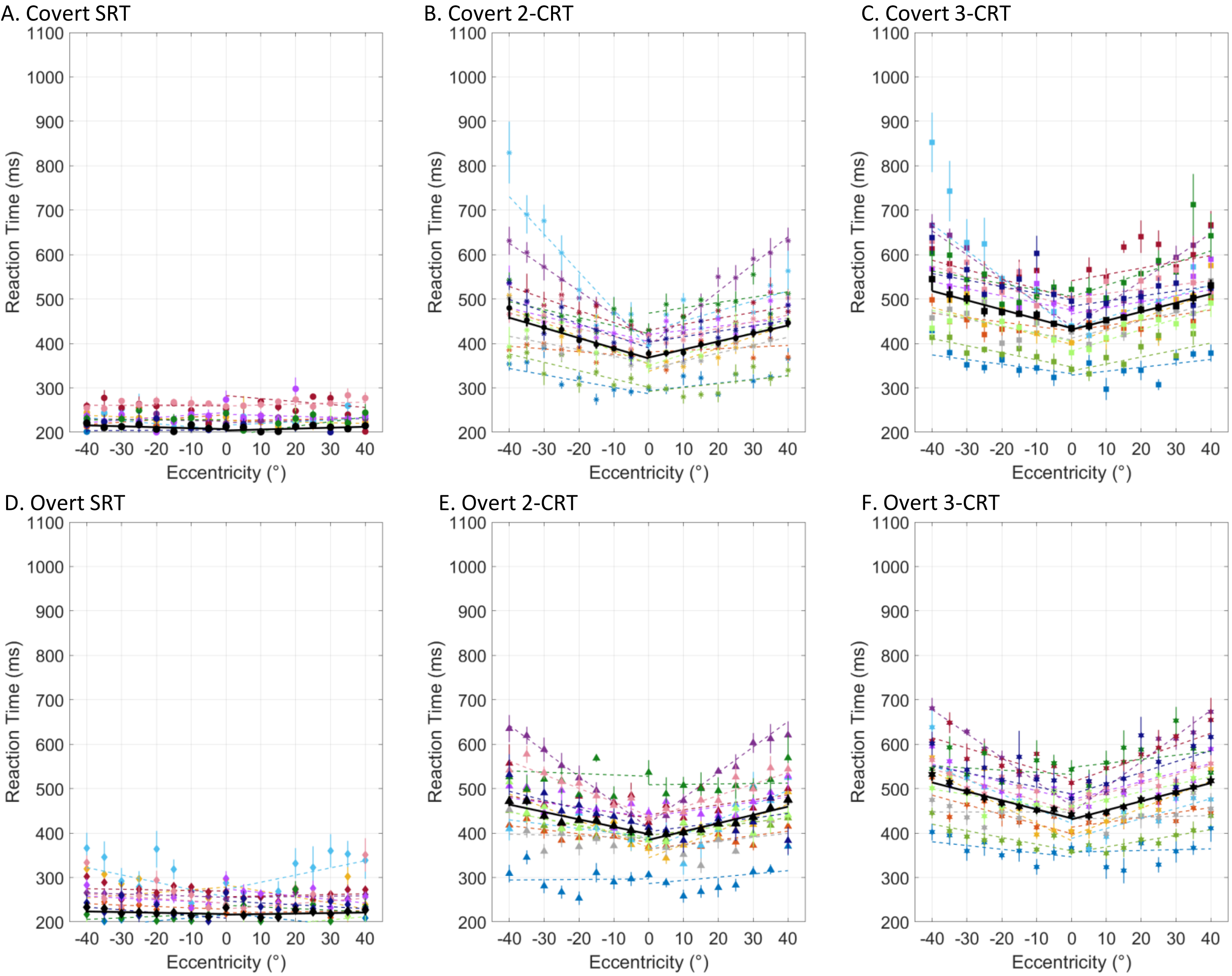


### Experiment 1a - Gabor

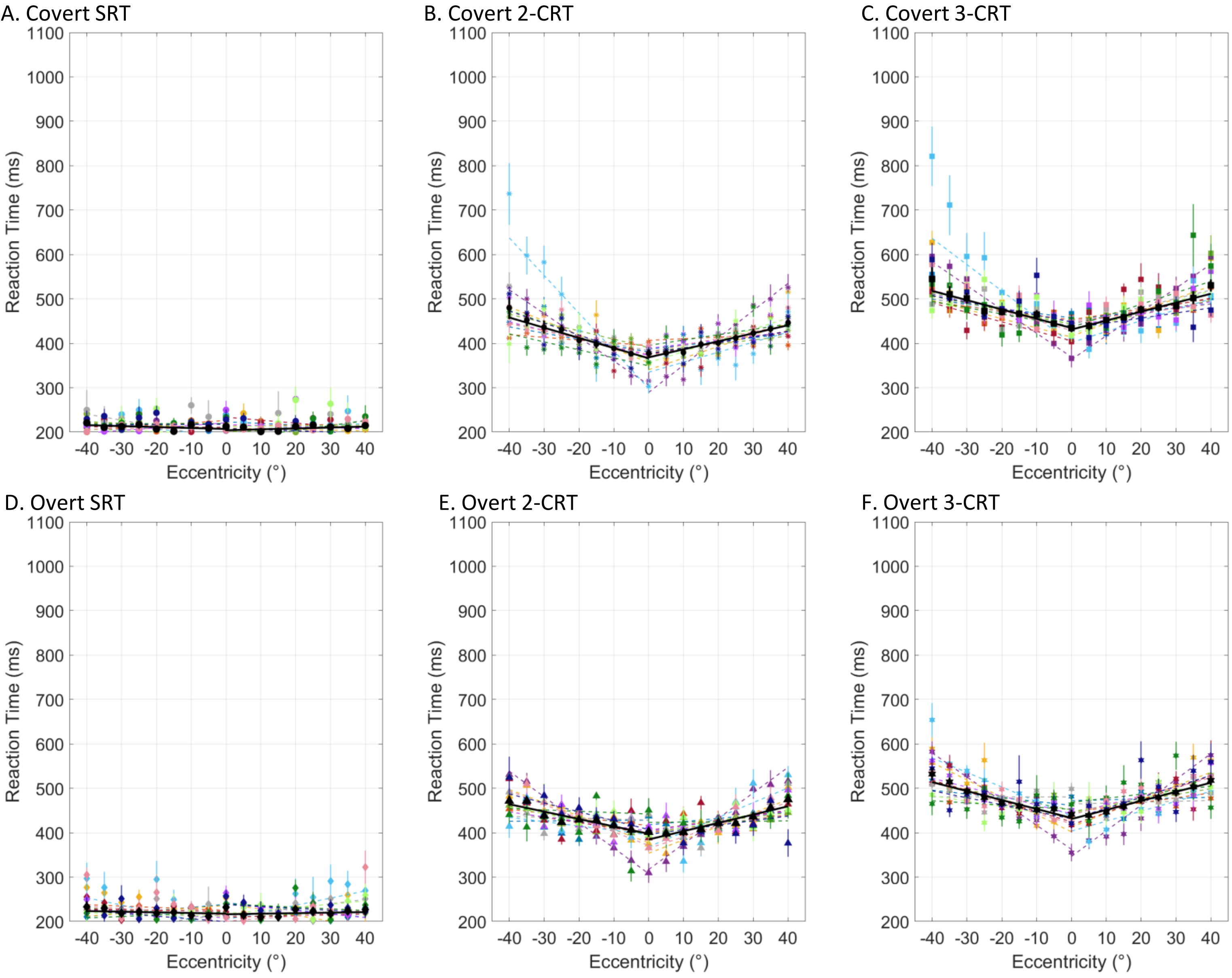


Experiment 1a – Gabor normalized

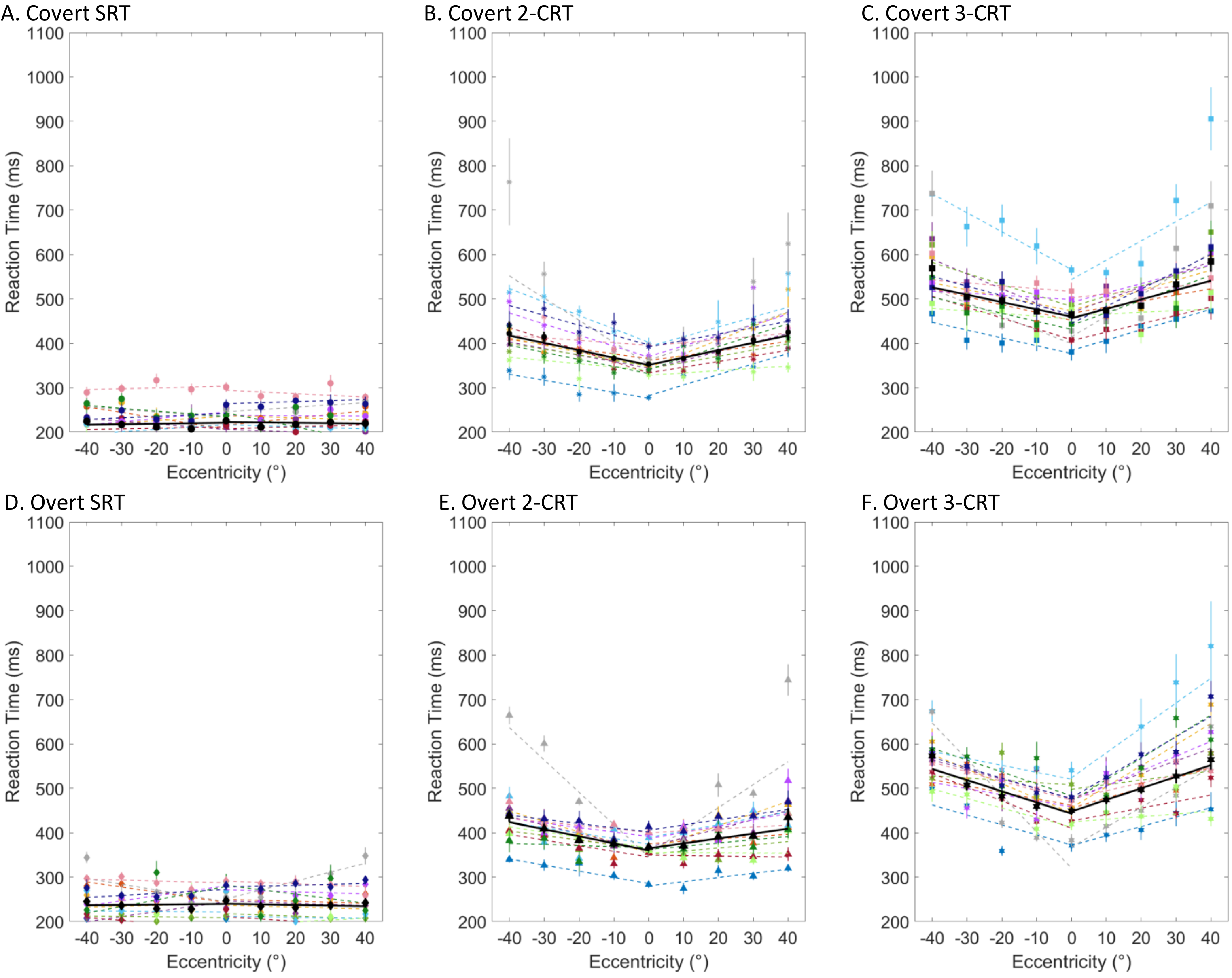


### Experiment 1b – Letter

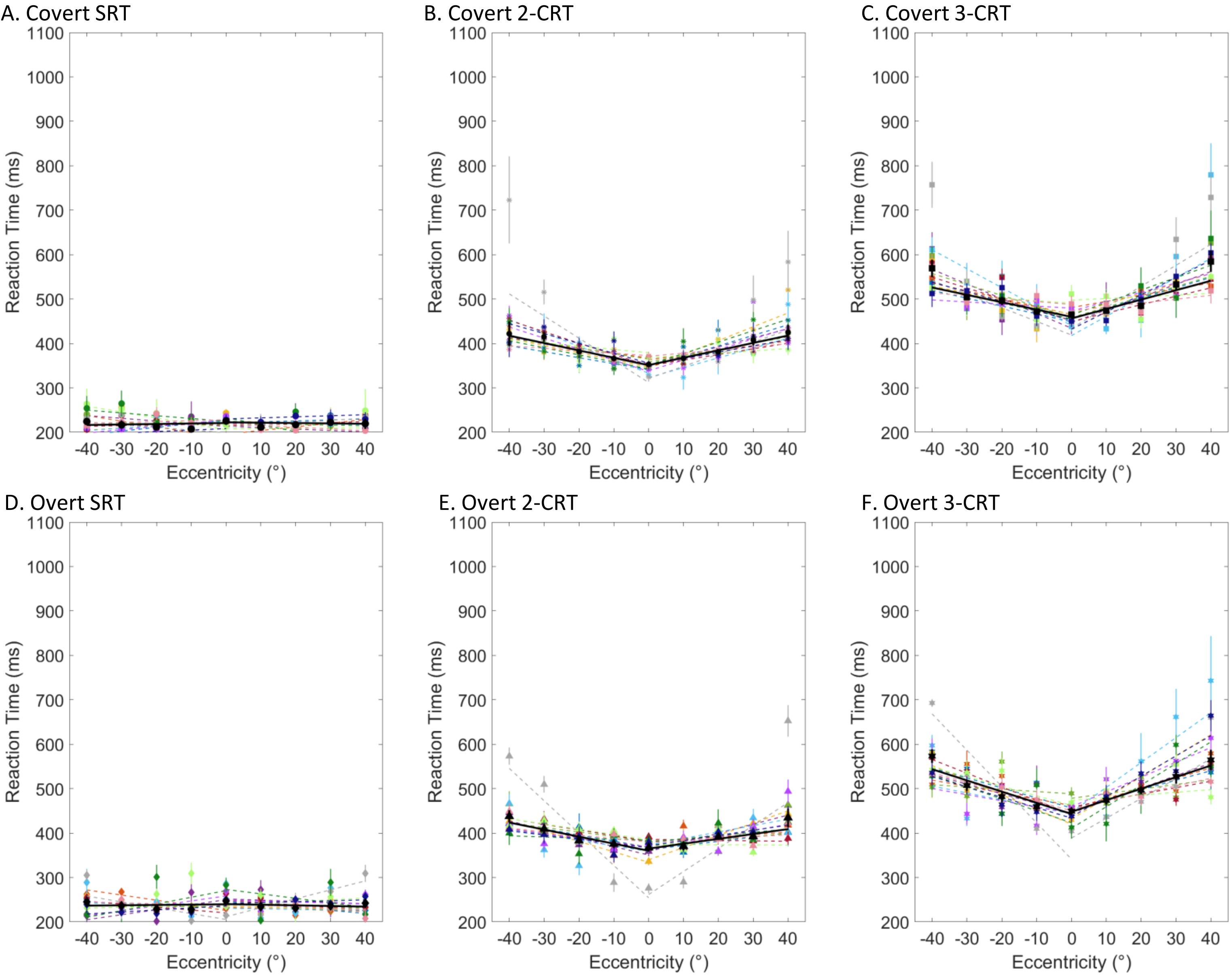


#### Experiment 1b – Letter normalized

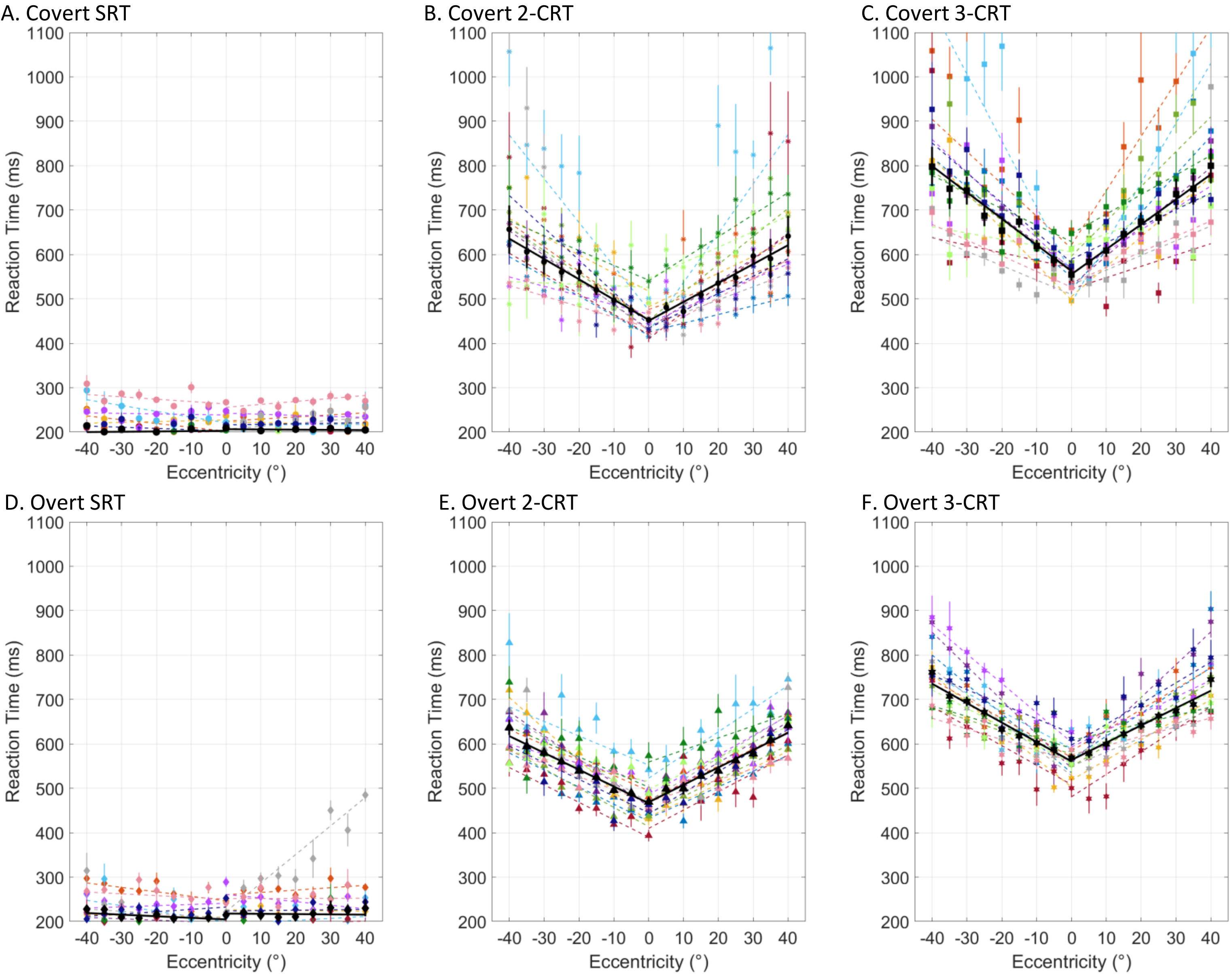


### Experiment 1c – Face

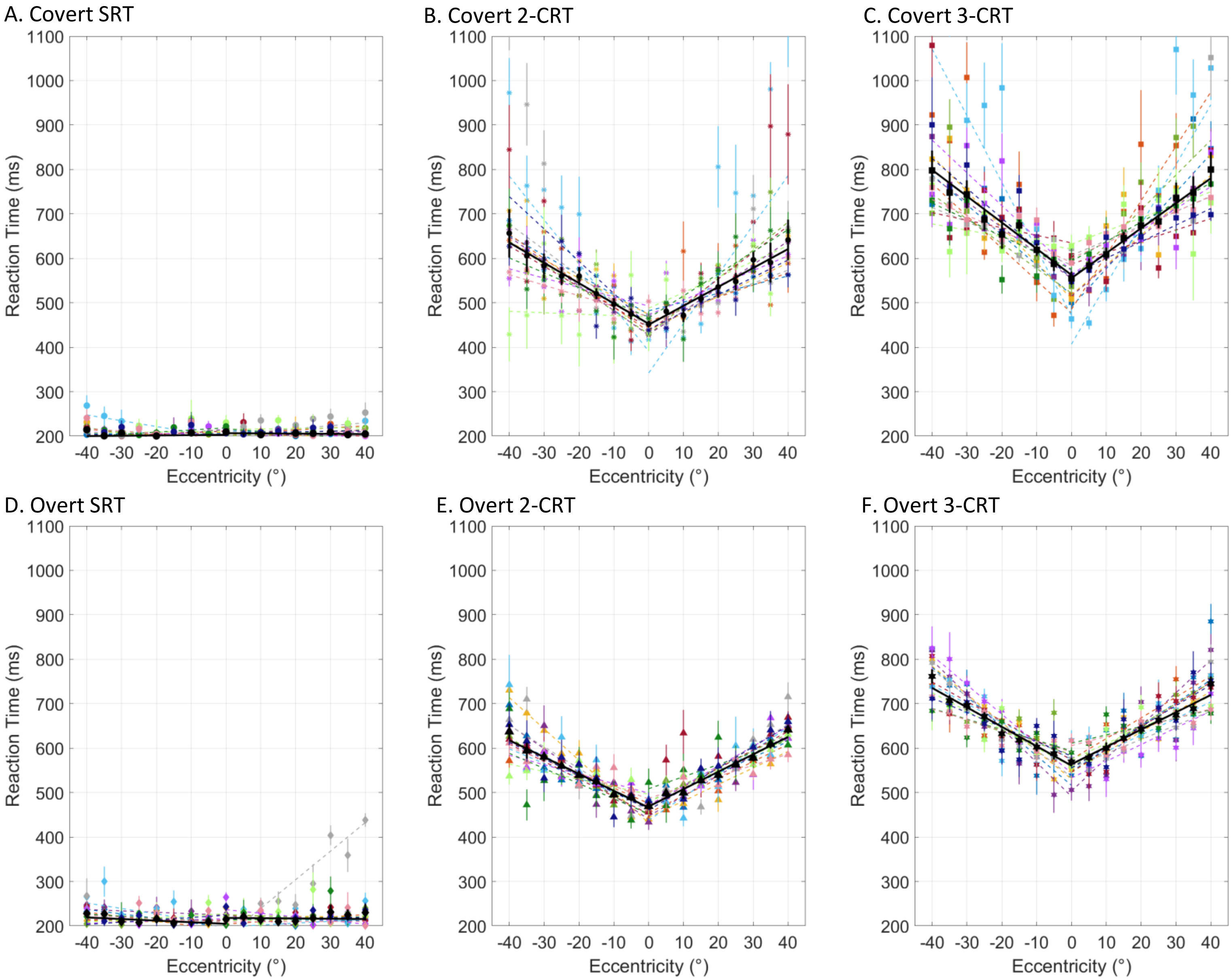


Experiment 1c – Face(a) normalized

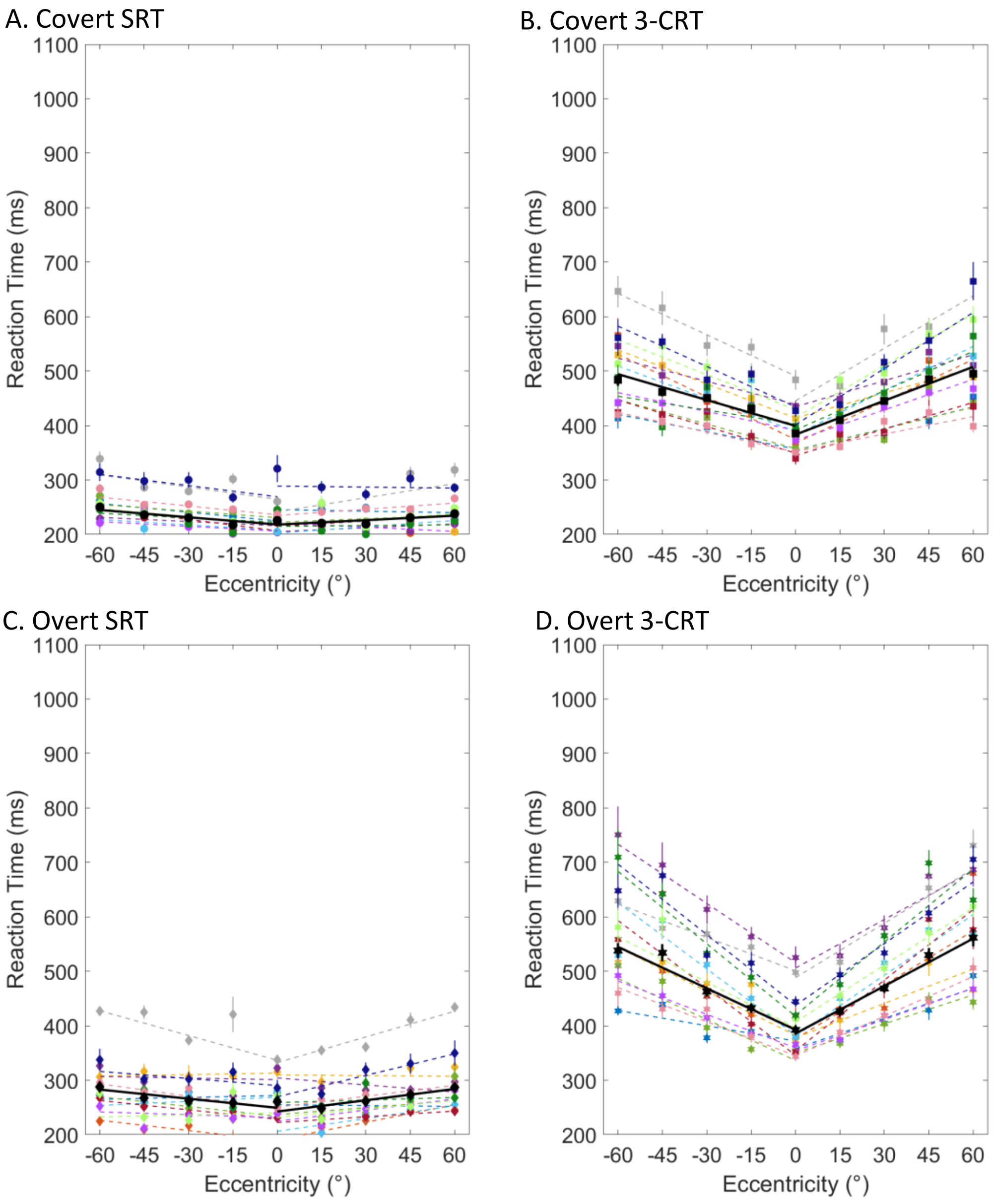


### Experiment 2a – BBB-H

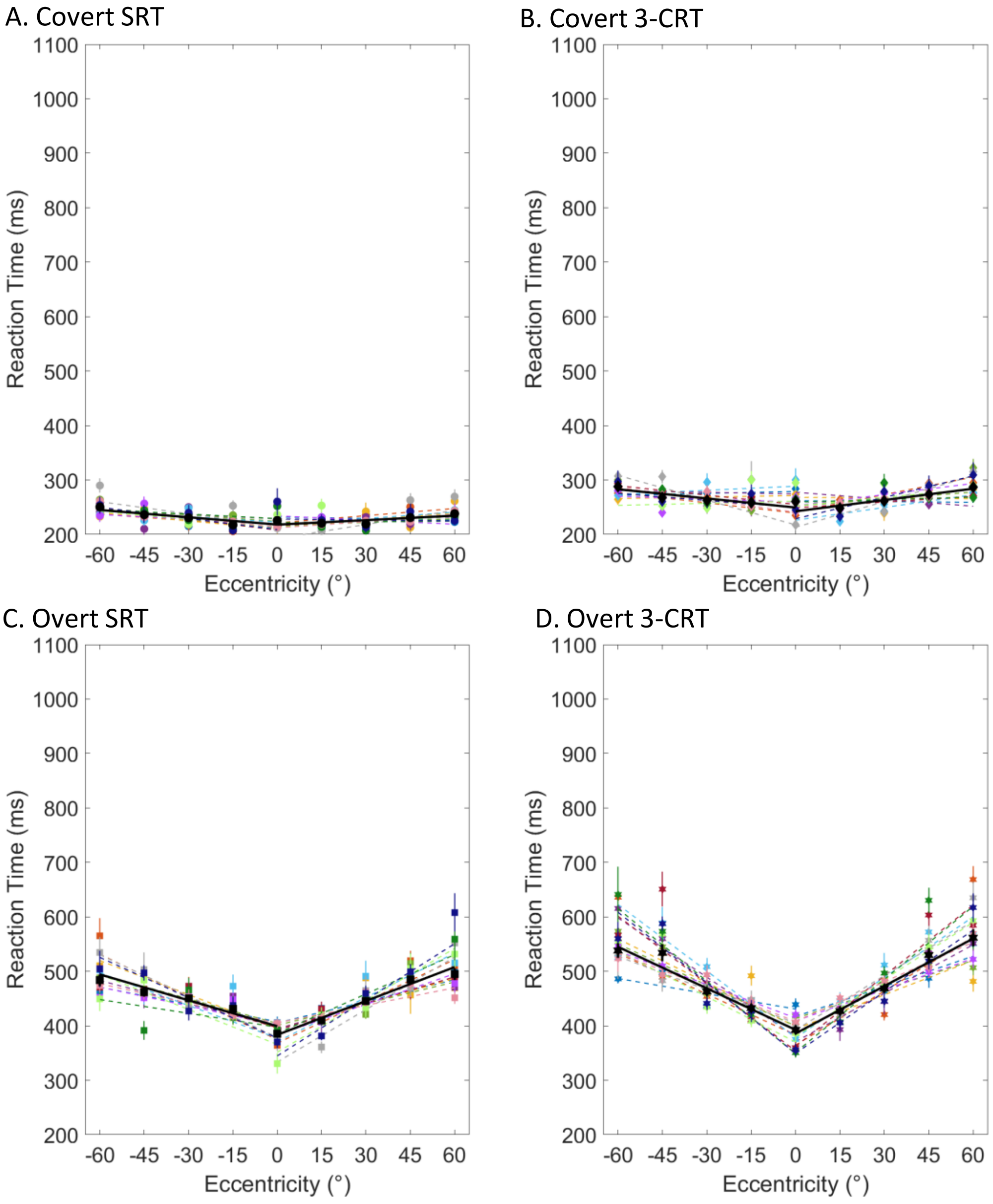


### Experiment 2a – BBB-H normalized

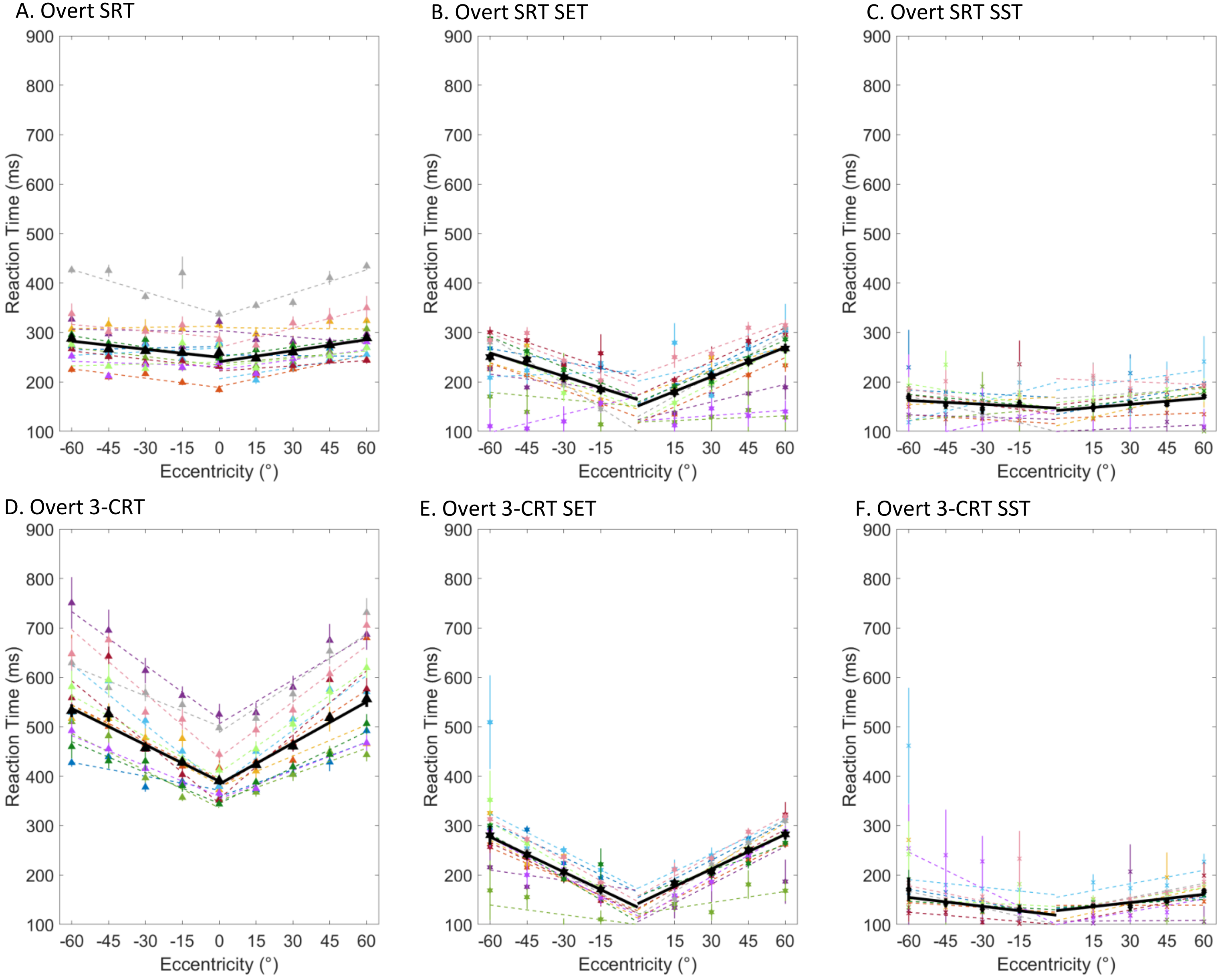


### Experiment 2a – BBB-H eye tracking

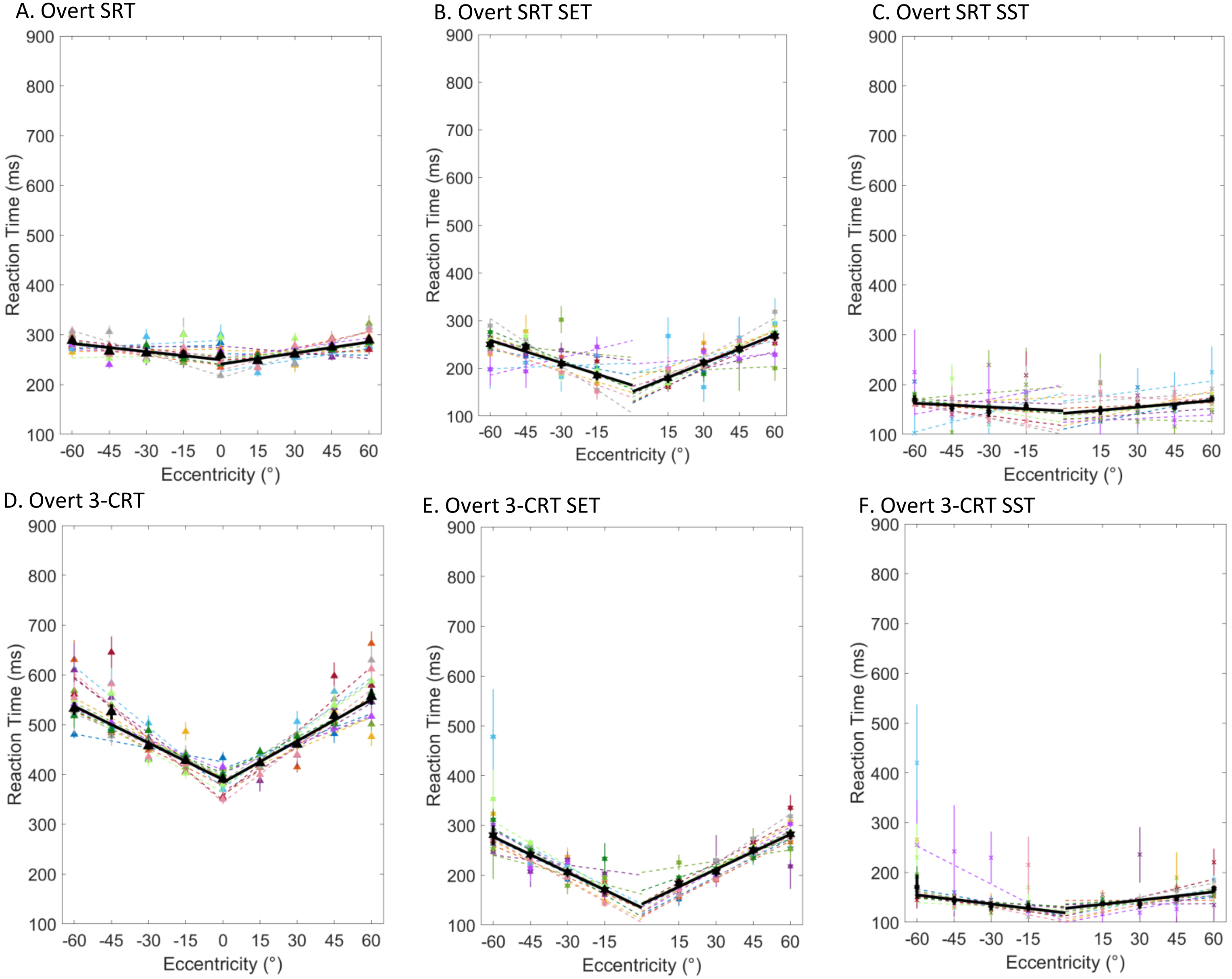


Experiment 2a – BBB-H eye tracking normalized

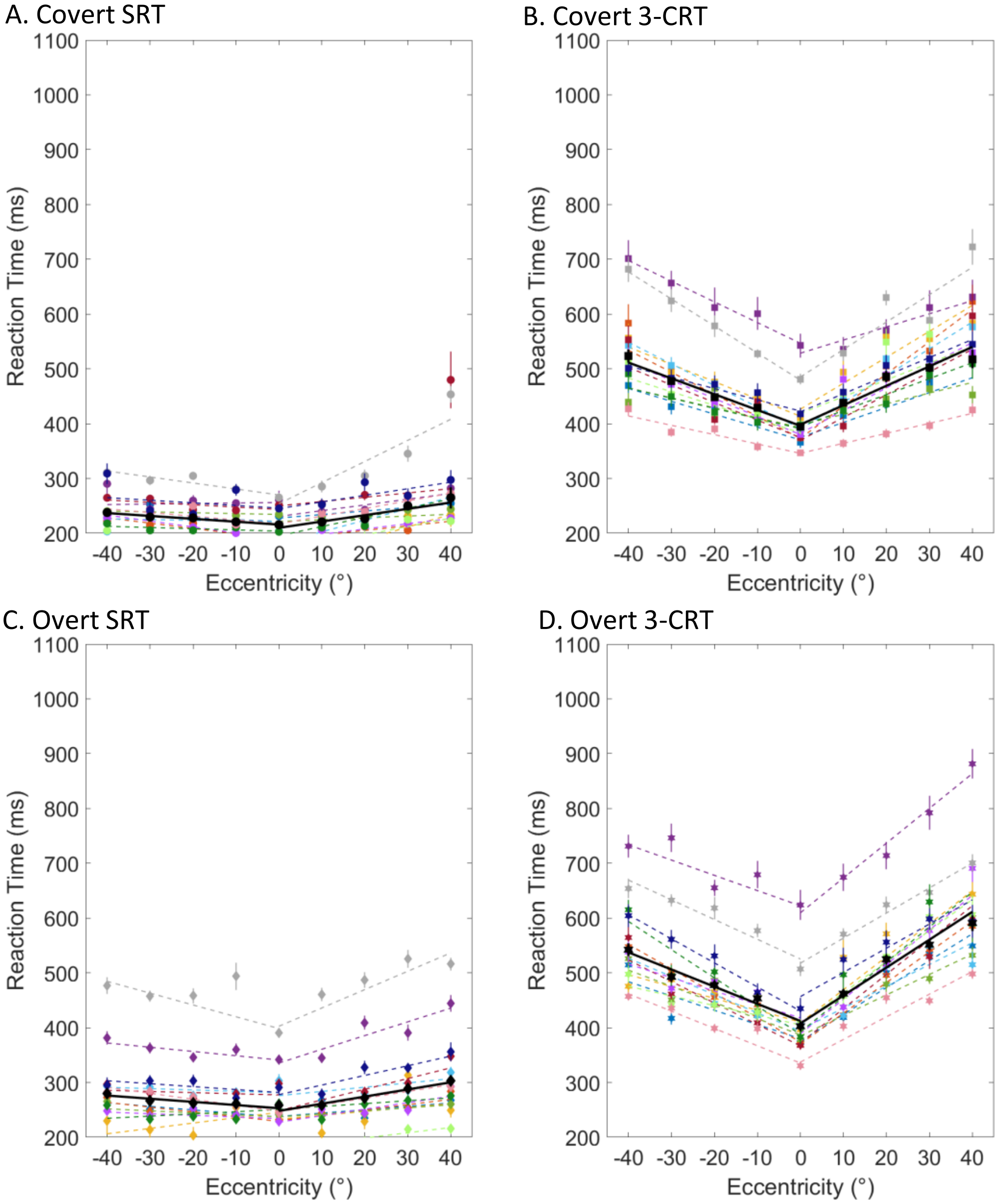


Experiment 2b – BBB-V

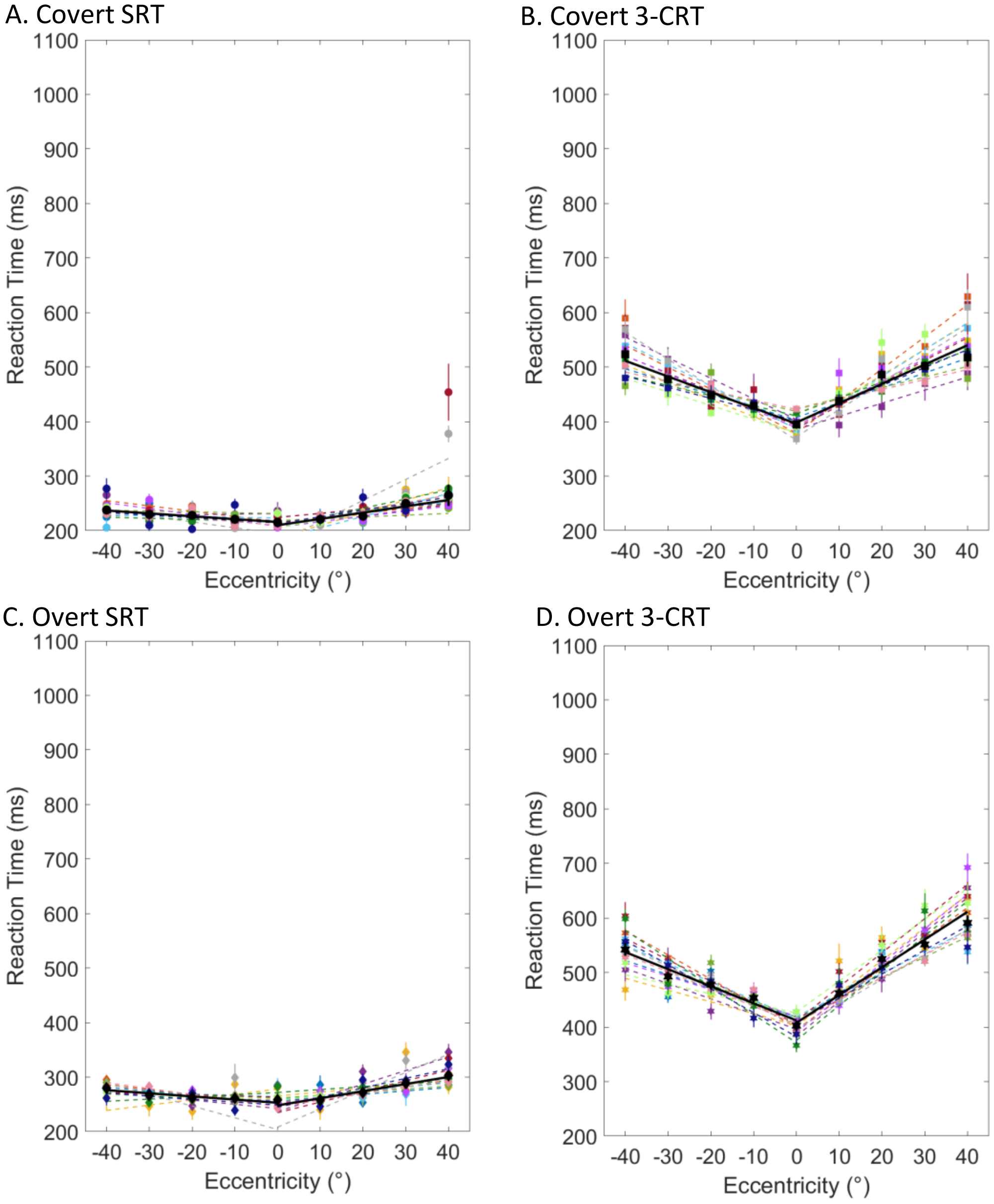


### Experiment 2b – BBB-V normalized

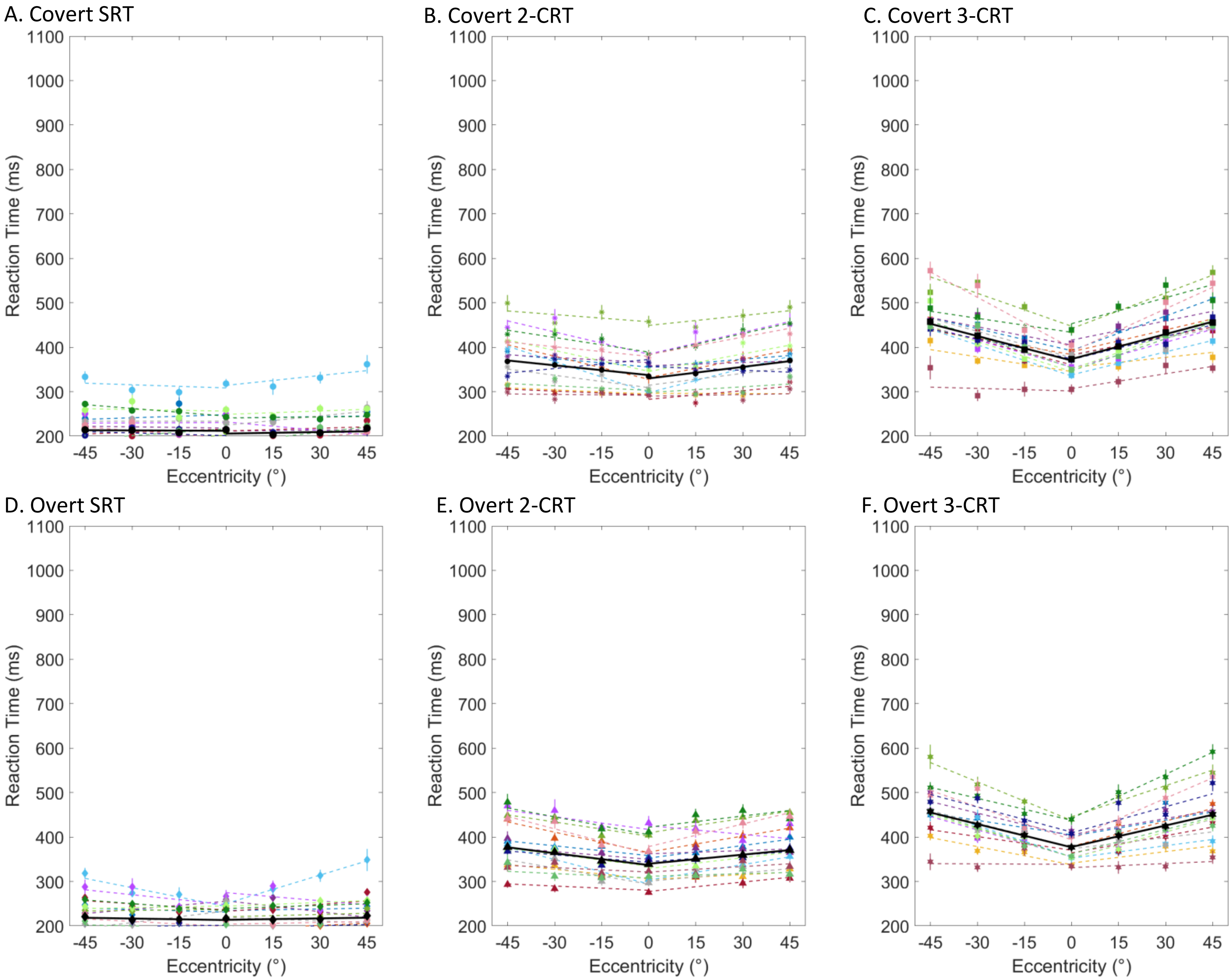


### Experiment 3a – BBB

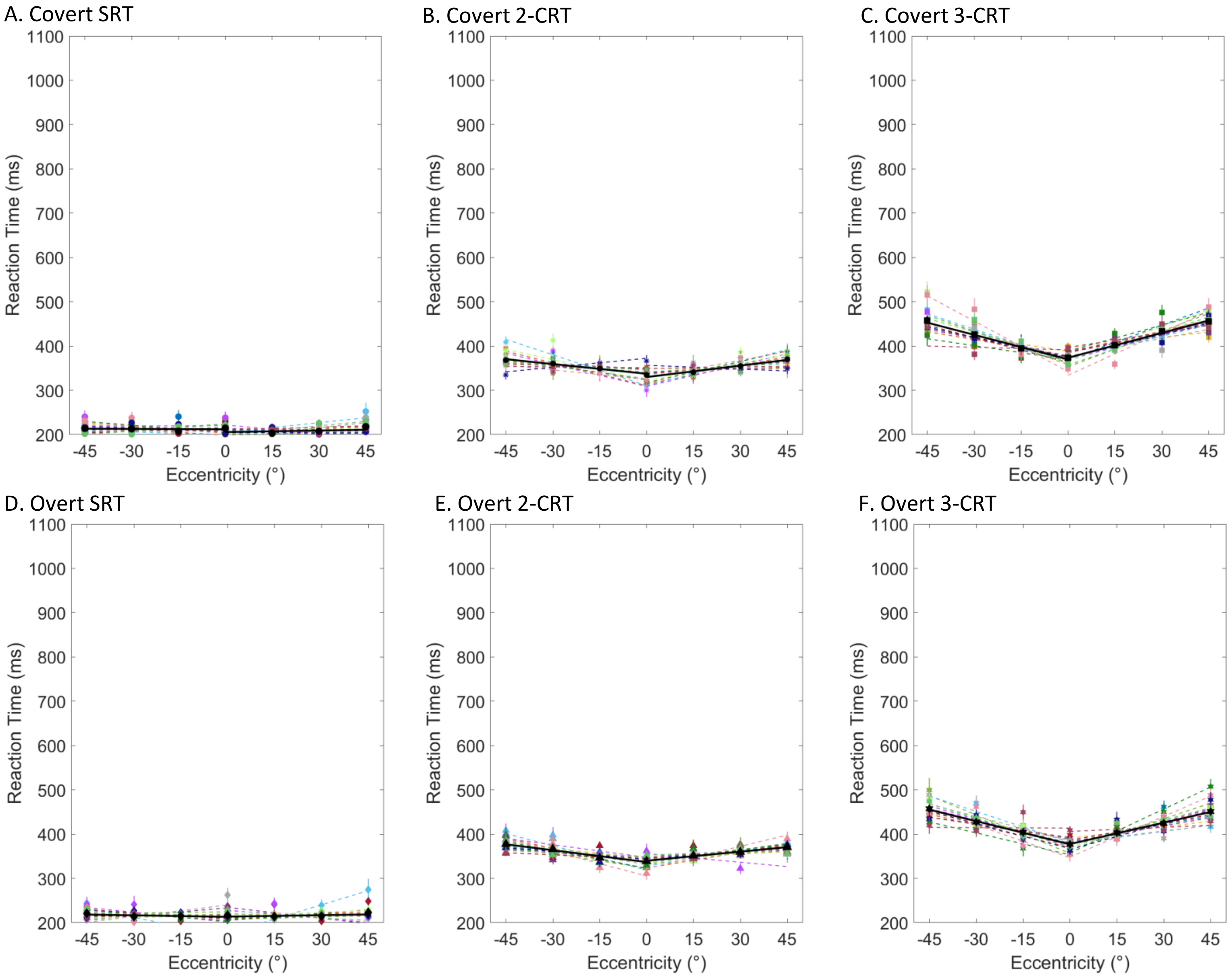


### Experiment 3a – BBB normalized

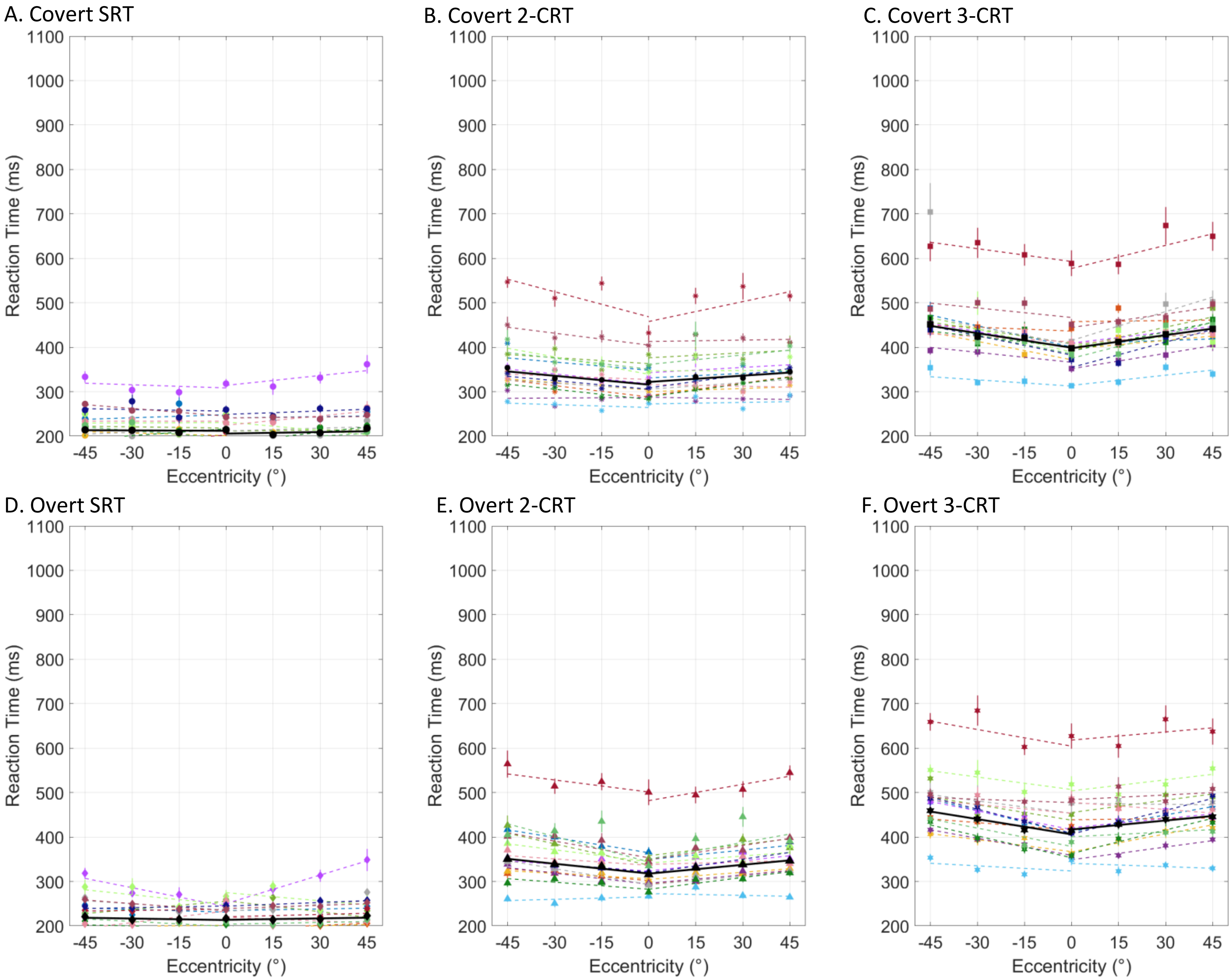


### Experiment 3b – Color

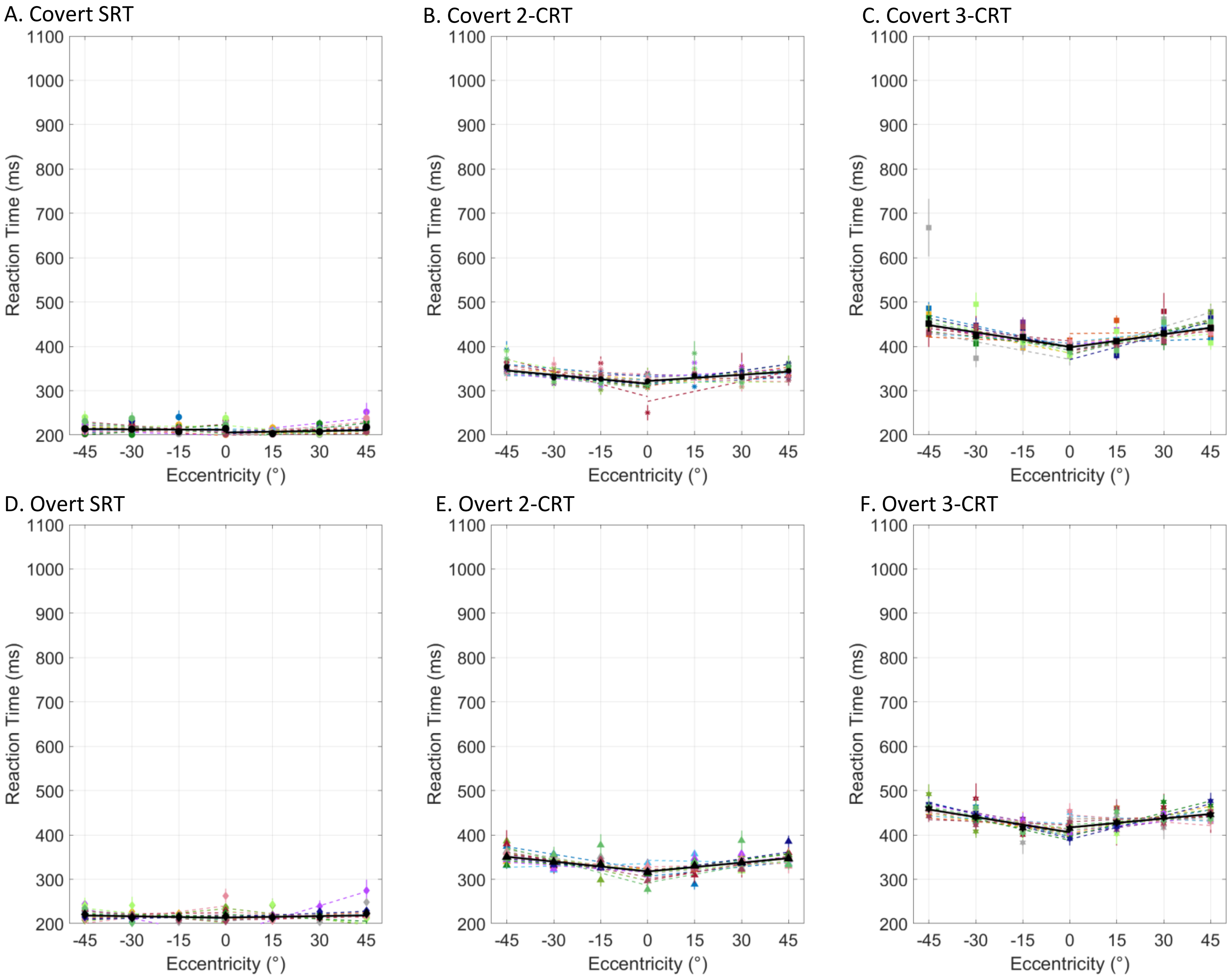


### Experiment 3b – Color normalized

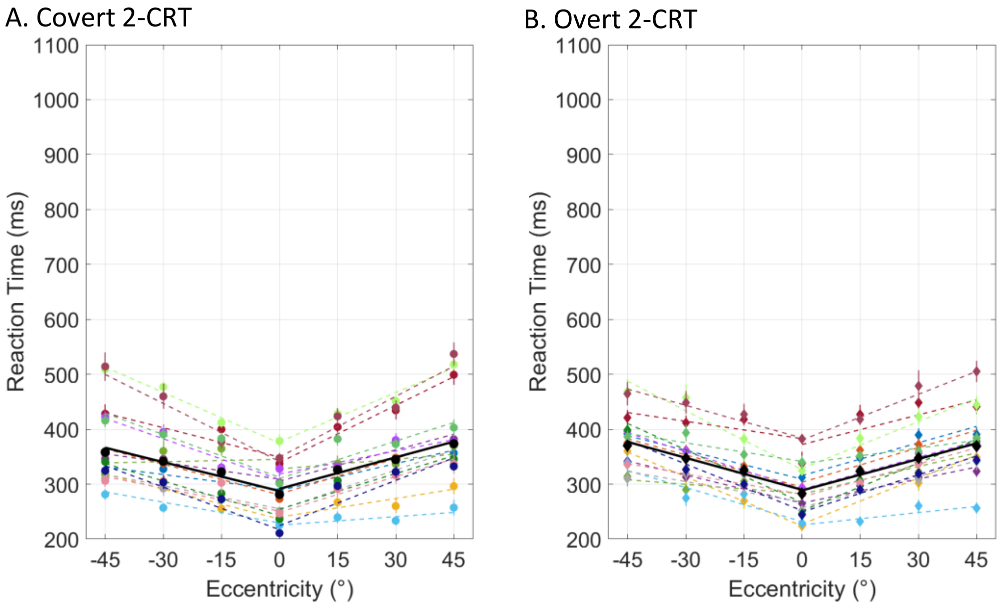


Experiment 3c – LR

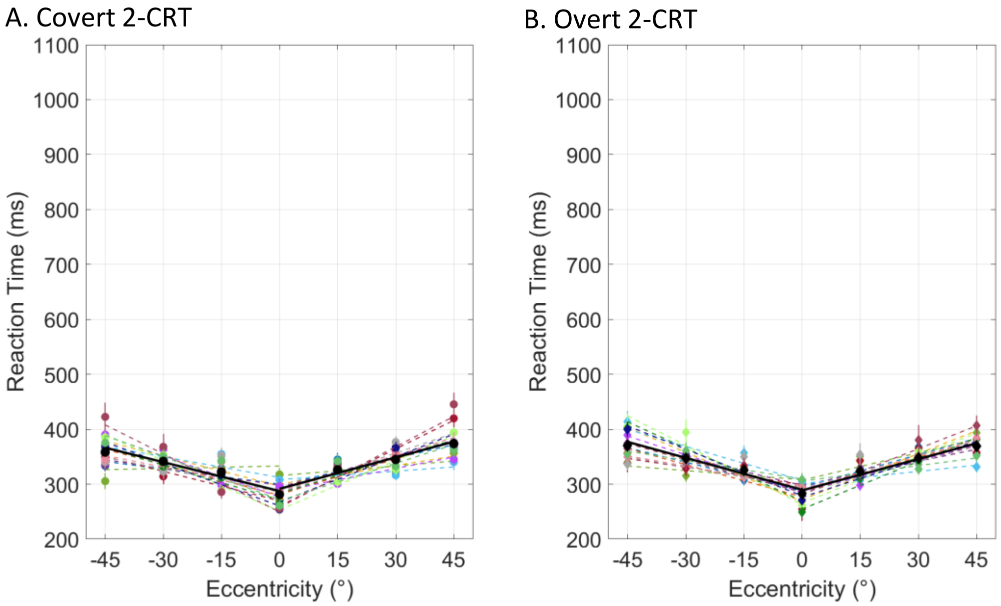


### Experiment 3c – LR normalized

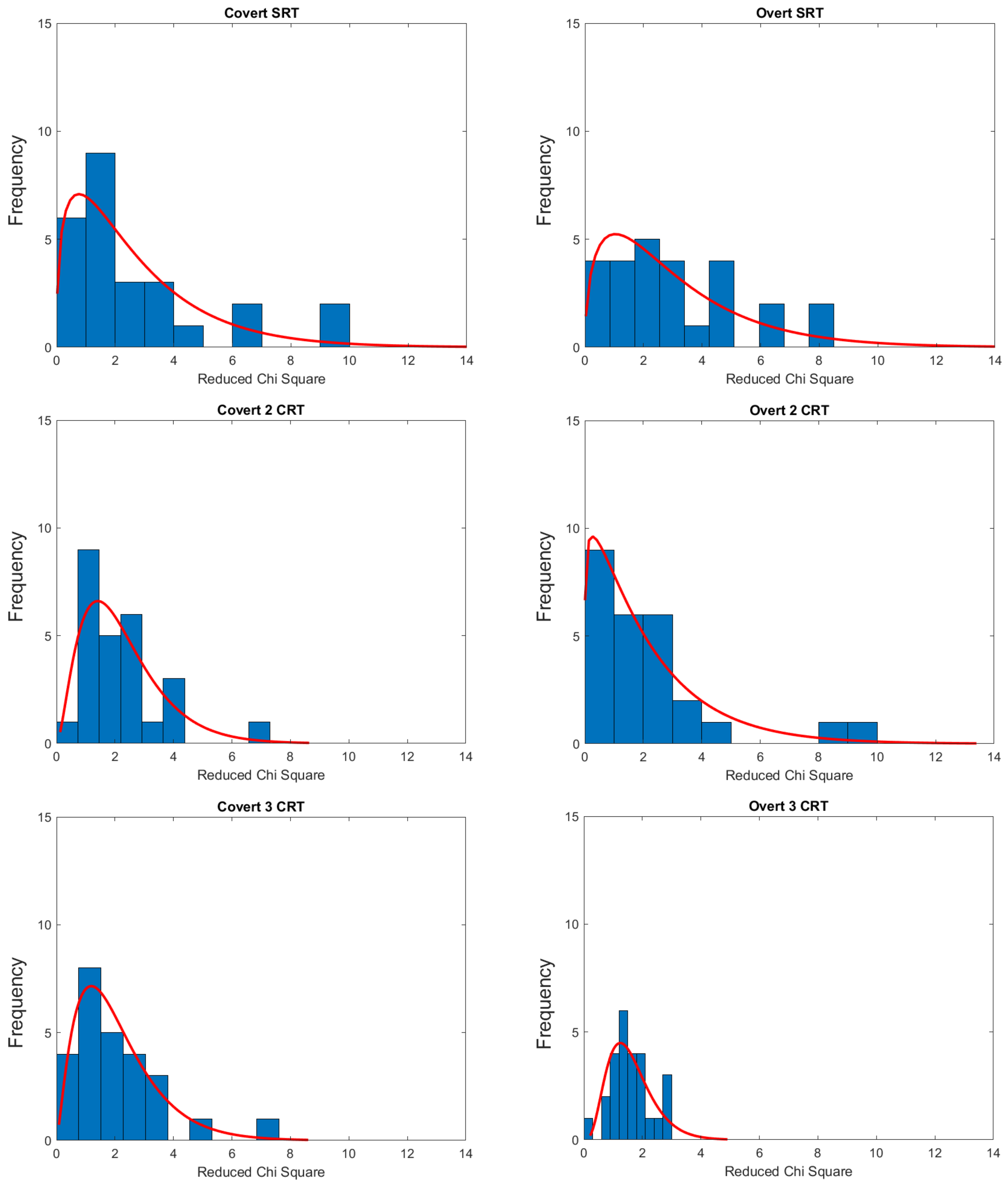


Experiment 1a - Gabor

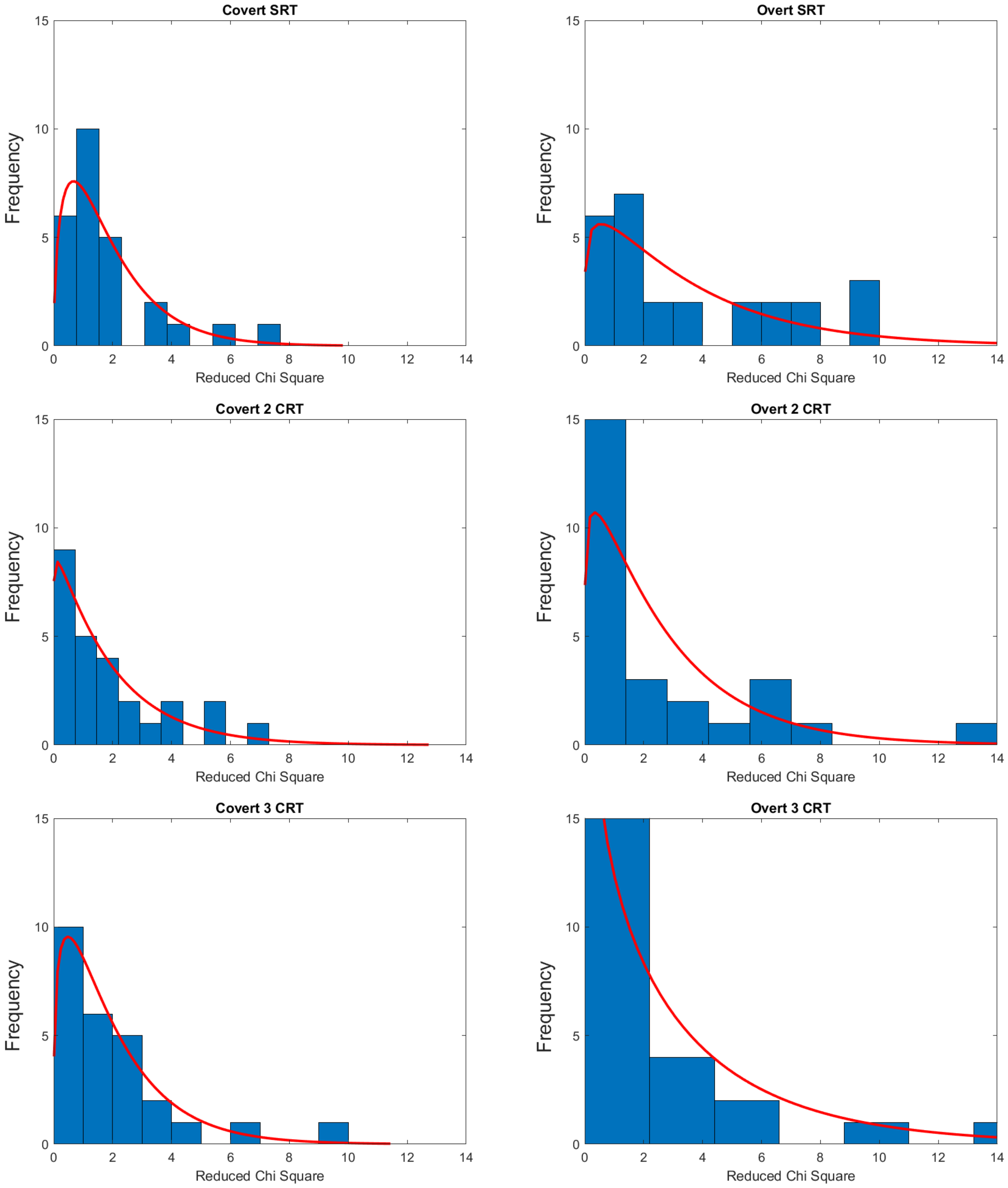


Experiment 1b - Letter

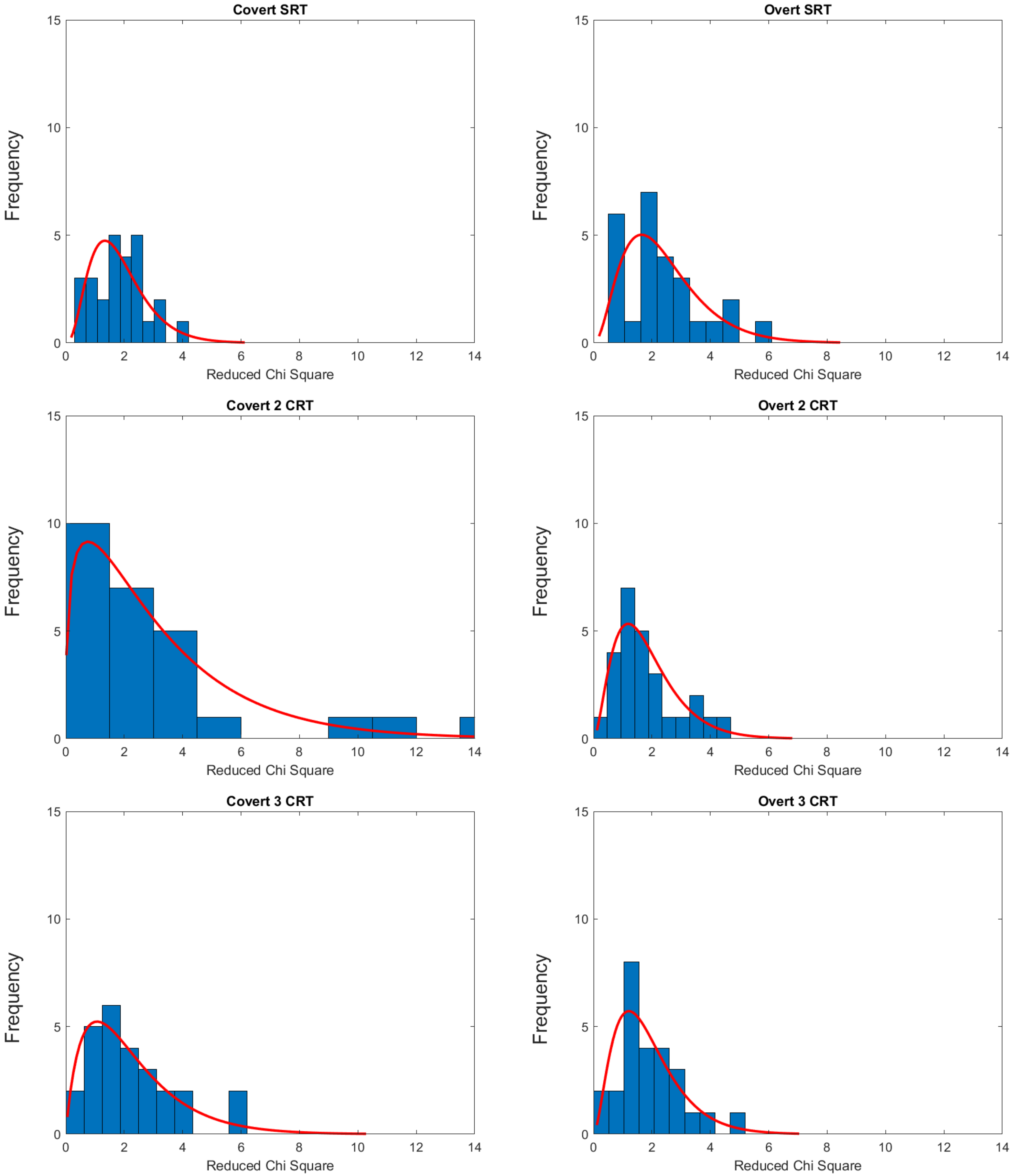


Experiment 1c – Face(a)

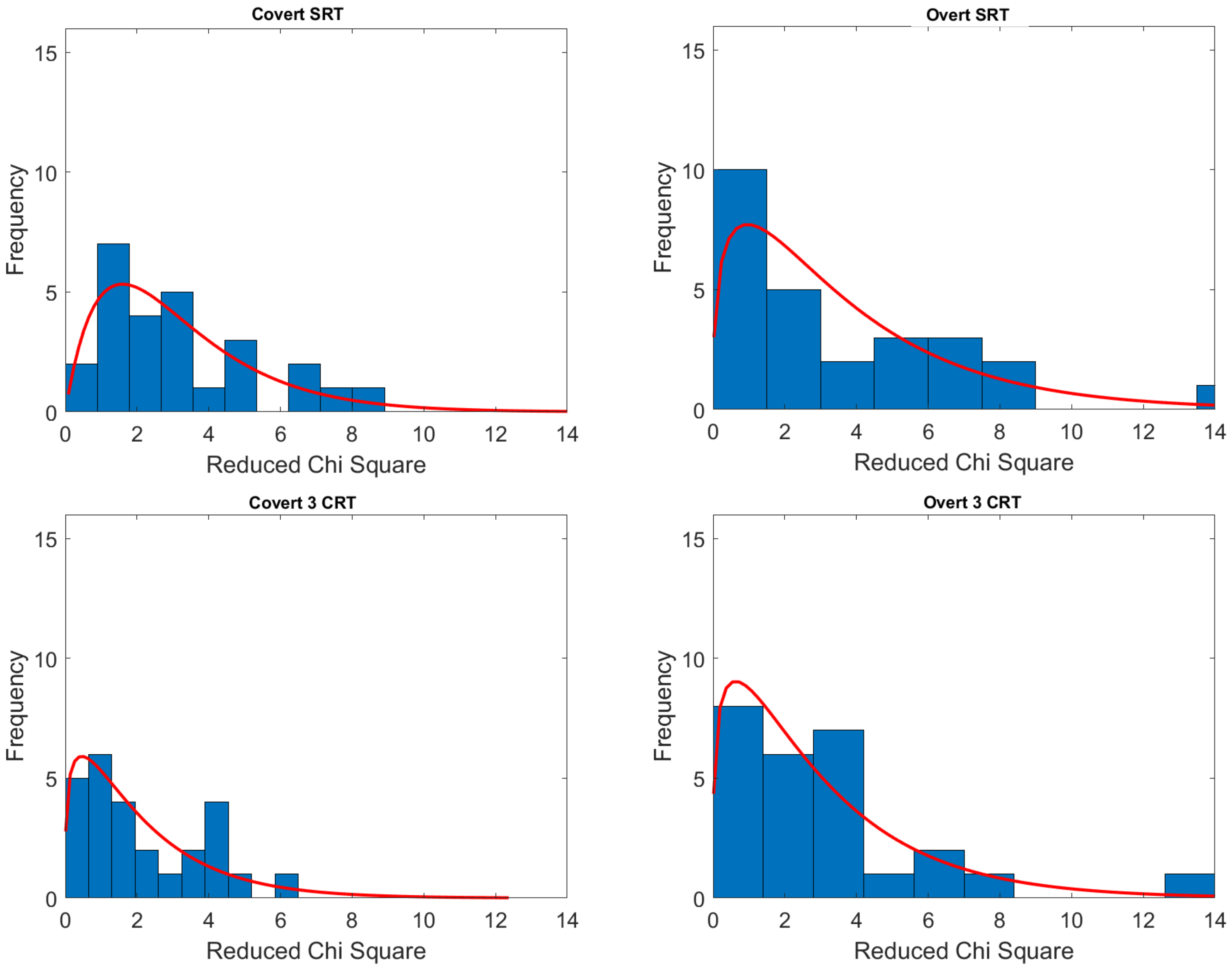


Experiment 2a – BBB-H

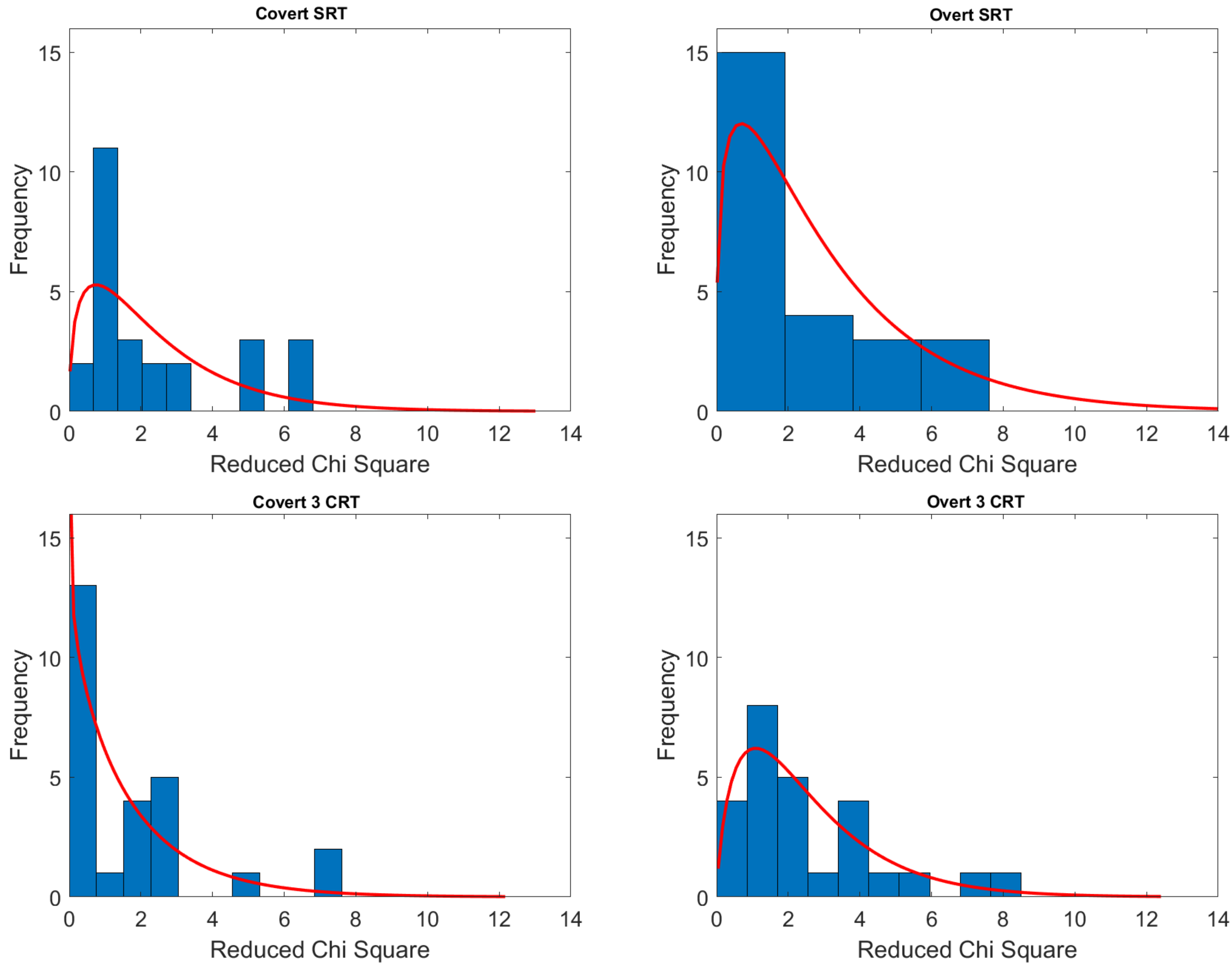


Experiment 2b – BBB-V

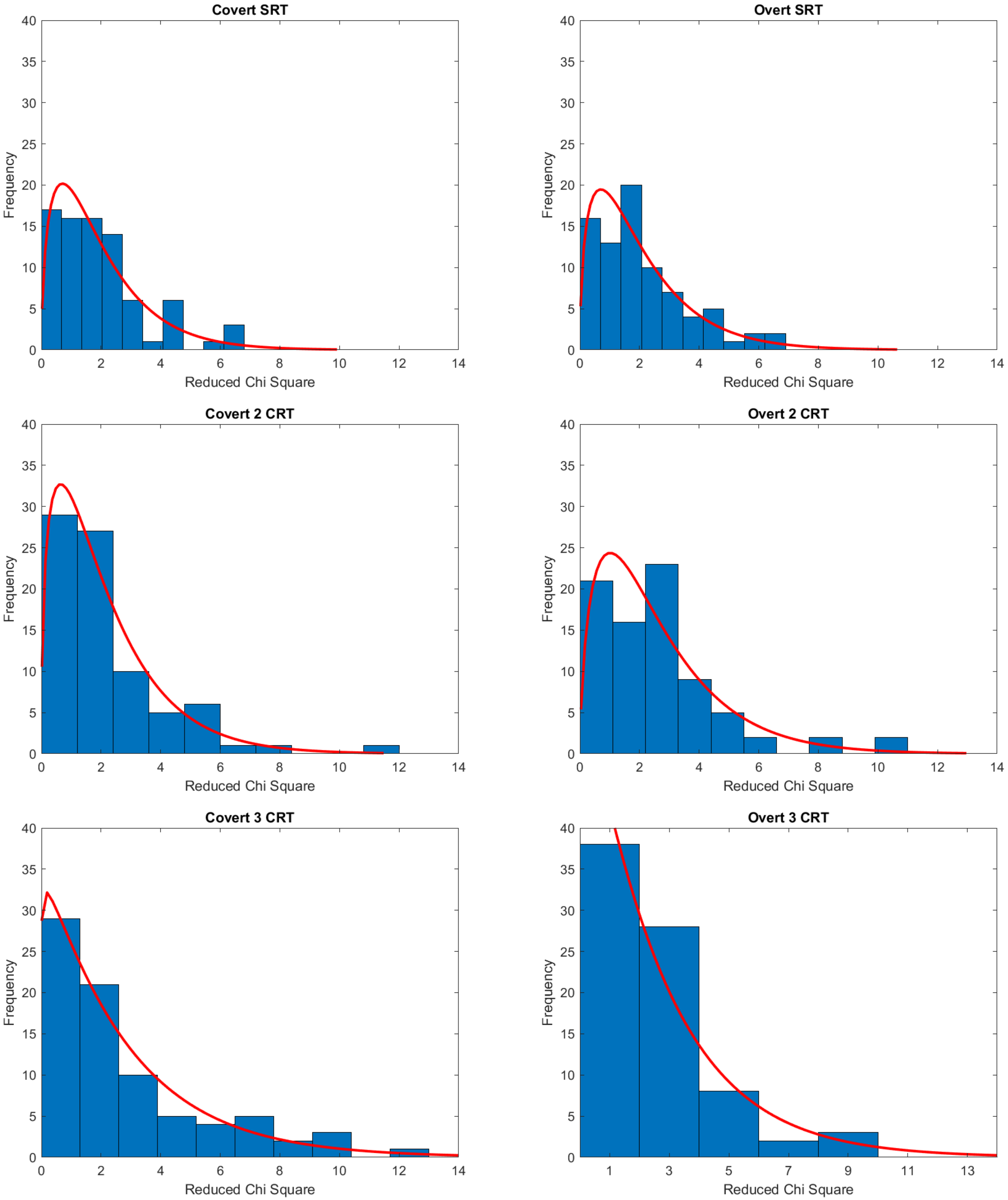


Experiment 2c – Face(b)

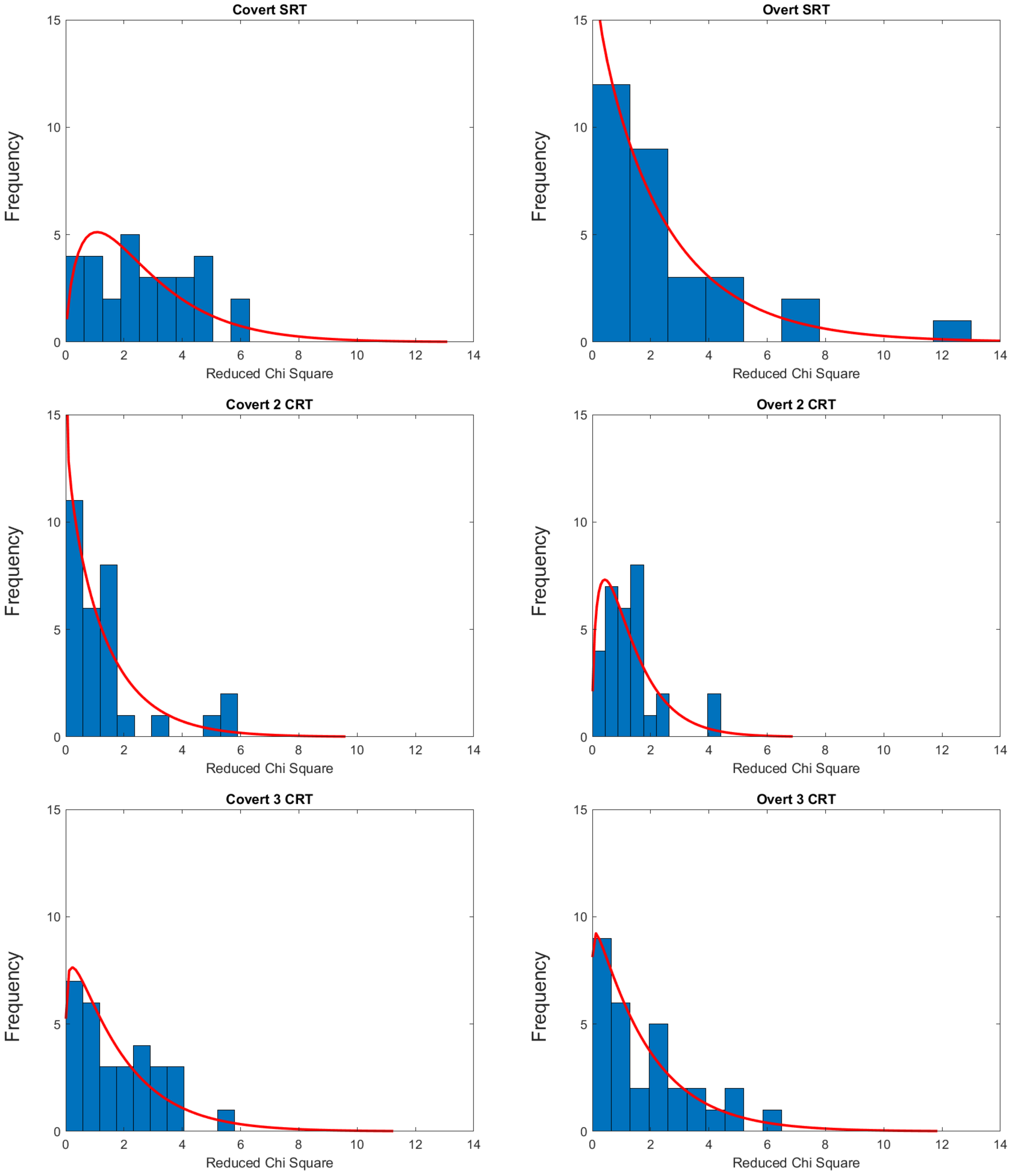


Experiment 3a – BBB

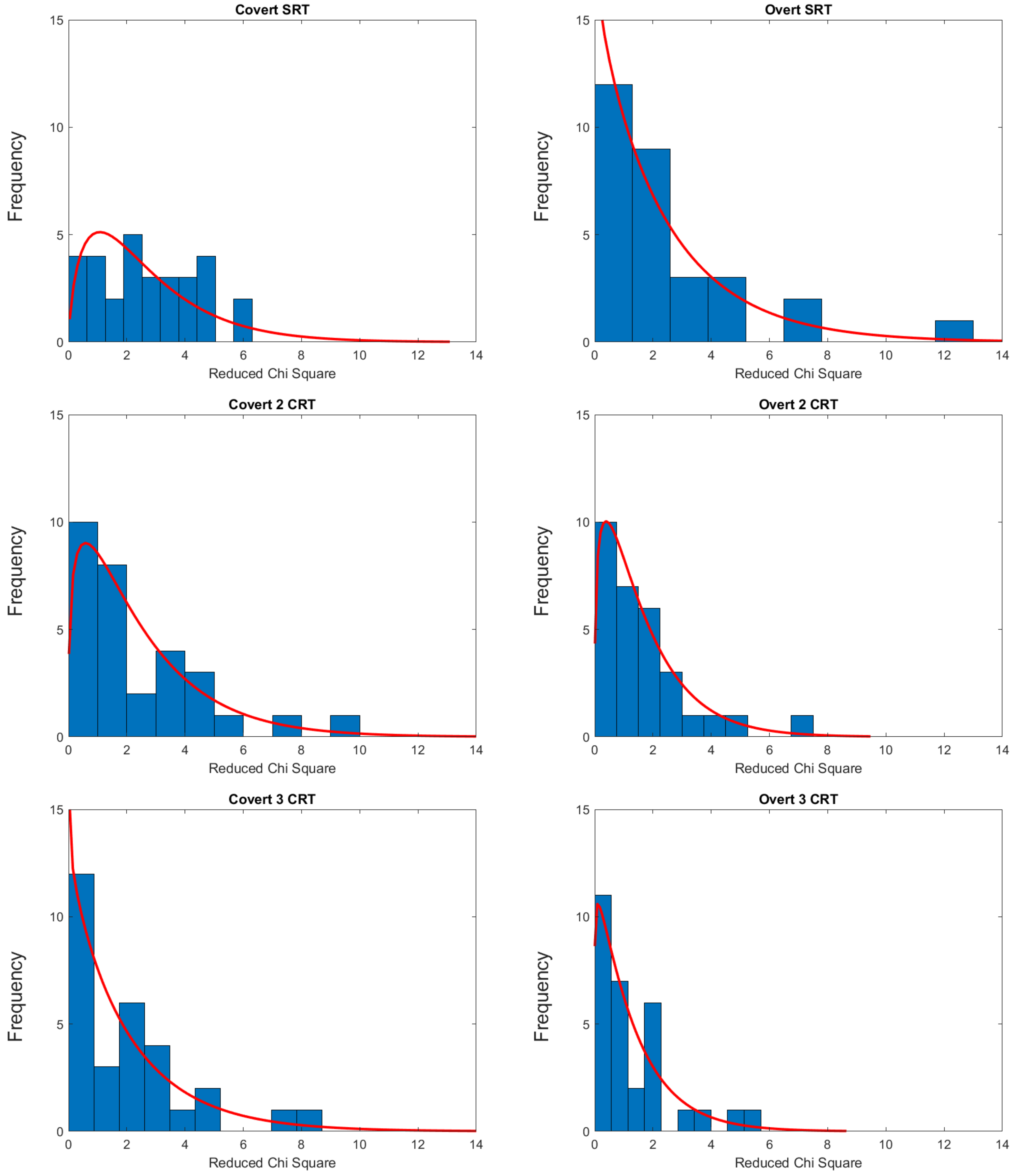


#### Experiment 3b – Color

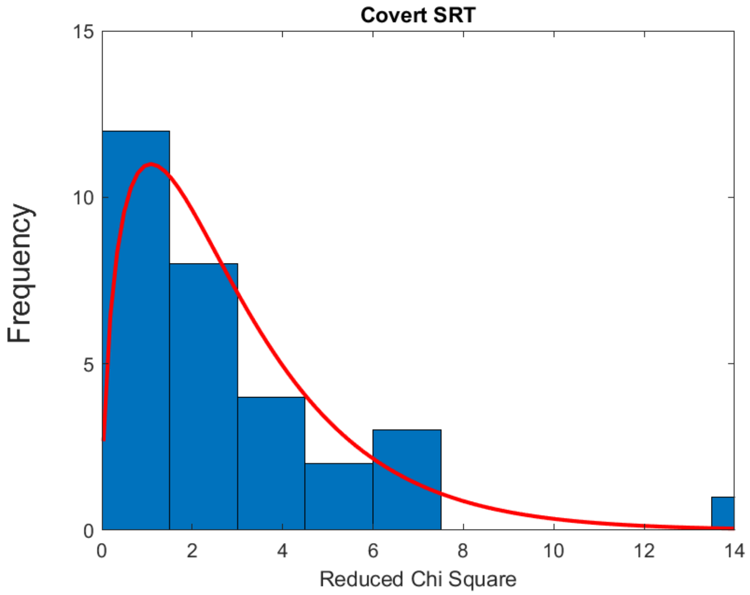


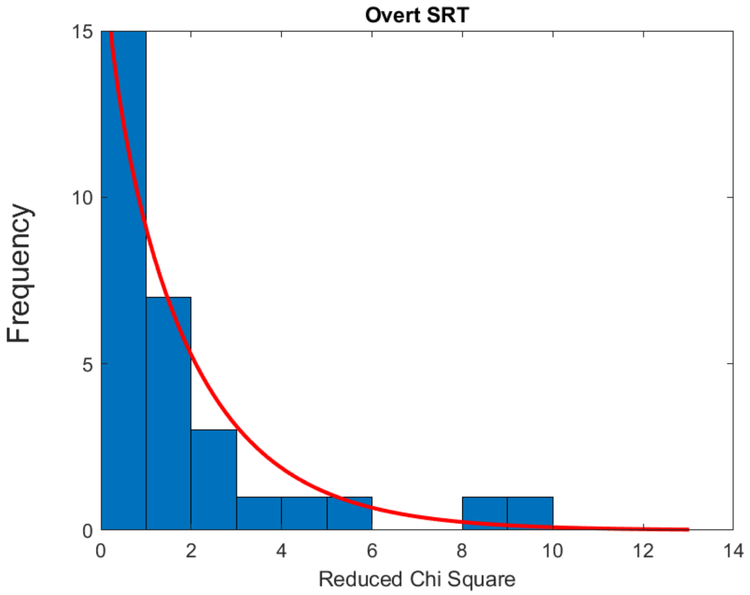


Experiment 3c – LR
